# Genetic Reinstatement of RIG-I in Chickens Reveals Insights into Avian Immune Evolution and Influenza Interaction

**DOI:** 10.1101/2023.11.01.564710

**Authors:** Hicham Sid, Theresa von Heyl, Sabrina Schleibinger, Romina Klinger, Leah Heymelot Nabel, Hanna Vikkula, Rodrigo Guabiraba, Vanaique Guillory, Ryan Scicluna, Mohanned Naif Alhussien, Brigitte Böhm, Benjamin Schade, Daniel Elleder, Samantha Sives, Lonneke Vervelde, Sascha Trapp, Benjamin Schusser

## Abstract

Retinoic acid-inducible gene I (*RIG-I*) activates mitochondrial antiviral signaling proteins, initiating the antiviral response. *RIG-I* and *RNF135*, a ubiquitin ligase regulator, are missing in domestic chickens but conserved in mallard ducks. The chickens’ *RIG-I* loss was long believed to be linked to increased avian influenza susceptibility. We reinstated both genes in chickens and examined their susceptibility to infection with an H7N1 avian influenza virus. Uninfected *RIG-I*-expressing chickens exhibited shifts in T and B cells. At the same time, the H7N1 infection led to severe disease, persistent weight loss, and increased viral replication compared to wild-type chickens. The simultaneous expression of *RIG-I* and *RNF135* potentiated the *RIG-*I activity and was associated with exacerbated inflammatory response and increased mortality without influencing virus replication. Additional animal infection experiments with two other avian influenza viruses validated these findings. They confirmed that the harmful effects triggered by *RIG-I* or *RIG-I*-*RNF135*-expression require a minimum degree of viral virulence. Our data indicate that the loss of *RIG-I* in chickens has likely evolved to counteract deleterious inflammation caused by viral infection and highlight an outcome of restoring evolutionary lost genes in birds.

**Significance Statement:** The evolutionary loss of a crucial innate immune sensor like *RIG-I* in domestic chickens and its presence in closely related avian species such as ducks has long puzzled researchers. We genetically reinstated *RIG-I* in chickens, alongside its ubiquitination factor, *RNF135*, to uncover their roles in responding to influenza virus interactions in chickens. Our findings suggest that the loss of *RIG-I* in chickens may have occurred as an adaptive strategy to mitigate harmful inflammation associated with influenza infection. We shed light on an outcome of reinstating evolutionarily lost genes in birds and open new avenues for understanding immune responses in vertebrates.

## Introduction

Avian influenza virus (AIV) is an epizootic pathogen with zoonotic potential (1) that recently caused devastating outbreaks worldwide, leading to the loss of millions of birds due to animal death and culling (2). The ability of the virus to spread between mammals and humans is highly concerning due to the potential risk of pandemics (3, 4). The viral reservoir of AIVs are wild birds of the orders *Anseriformes* (ducks, geese, and swans) and *Charadriiformes* (gulls and terns) (5), which, compared to chickens or other galliform birds, exhibit milder clinical symptoms despite efficient viral replication (6). Certain genomic features of the duck, including a functional retinoic acid-inducible gene I *(RIG-I)* gene, were reported to be associated with their relative resistance to clinical avian influenza infection (7). *RIG-I* is a cytosolic RNA sensor that recognizes and binds to the 5’ triphosphate end (5’-ppp) (8). It forms a first line of antiviral defense as a pathogen recognition receptor (PRR) against RNA viruses (9). Upon activation, the *RIG-I* interacts with the mitochondrial antiviral signaling proteins (MAVS), inducing a pro-inflammatory antiviral response characterized by the upregulation of type I and type III interferons (IFNs) followed by the expression of IFN stimulated genes (ISGs) (10). The activity of *RIG-I* is believed to be controlled by post-translational modification of tripartite motif-containing protein 25 (*TRIM25*) (11) and RING finger protein 135 (*RNF135*, also known as *Riplet* or *REUL*). The latter was found to modify *RIG-I* by lysine 63-linked polyubiquitination of the C-terminal region of the caspase activation and recruitment domain (CARD) (12), leading to a stronger *RIG-I* signal transduction (13). The physiological role of mammalian *RIG-I*, its function in viral infections, and its parallel evolution with other RLRs are well documented (11, 14–16). Furthermore, the knockout (KO) of *RIG-I* in mice has shown that this gene may have additional functions related to adaptive immunity (14). *RIG-I* KO mice exhibited a defect in migratory dendritic cells in combination with a reduced frequency of polyfunctional effector T cells upon influenza infection (17).

In ducks, *RIG-I* elicits a potent interferon (IFN) response within the first few hours after infection, leading to survival against most AIV strains (18). In contrast, chickens lack *RIG-I*, which probably lost its function in a common ancestor of galliform birds (7, 19). Interestingly, a recent study detected disrupted *RIG-I* pseudogenes in some Galliformes, including the helmeted guineafowl (*N. meleagris*) and the northern bobwhite (*C. virginianus*) (20). Authors hypothesized a compensatory evolution of melanoma differentiation-associated gene-5 (*MDA5*) that accompanied the gradual loss of *RIG-I* in chickens (20). The evolutionary loss of *RIG-I* in different galliform birds correlated with the simultaneous loss of its ubiquitin ligase *RNF135* (20), which has been described to be critical for *RIG-I* ubiquitination in mammals, unlike *TRIM25*, which is increasingly believed to be less important for *RIG-I* signaling (21). Reasons behind the loss of *RIG-I* and its ubiquitination factor in chickens are still unknown and remain enigmatic, especially given the virus-limiting effect of duck *RIG-I* overexpression in AIV-infected chicken DF-1 cells (7).

To date, the effect of *in vivo* expression of *RIG-I* in chickens has never been studied, and no transgenic chicken lines expressing duck *RIG-I* have been generated, likely due to the lack of suitable biotechnological tools in avian research. Here, we used chicken primordial germ cells (PGCs) to develop genetically modified chickens expressing duck *RIG-I* and *RNF135* under the control of their respective duck promoters. The generated birds were healthy and developed normally compared to their WT siblings. In the absence of infection, we observed differences in adaptive immune cells of *RIG-I*-expressing chickens, particularly T cell populations. In contrast, the co-expression of *RNF135* with *RIG-I* contributed to a balanced adaptive immune phenotype that appeared to be similar to WT birds. Infection experiments with an H7N1 AIV led to severe clinical disease associated with a strong inflammatory response, high *IFN-γ* expression, and elevated viral replication in *RIG-I*-expressing chickens compared to other challenged groups. In contrast, infected *RIG-I-RNF135*-expressing chickens presented with inflammation and a differential expression of *IFN* and pro-inflammatory cytokines compared to *RIG-I*-expressing chickens. The obtained data reveal the immunological functions of *RIG-I* in chickens and the benefit of learning from less susceptible species to influenza infection to improve the immune system’s resilience towards infection.

## Results

### Generation of PGCs that express duck *RIG-I* and *RNF135* under the respective duck promoters

The genetic modification of PGCs represents a crucial step for generating transgenic chickens. PGCs are the precursors of sperm and eggs in adult animals, and therefore, we used them to produce chimeric roosters paired with WT layer hens to obtain the desirable transgene. To generate chickens that express *RIG-I* and *RNF135*, we started by cloning duck *RIG-I* and *RNF135*, which were subsequently inserted into two different expression vectors. *RIG-I* and *RNF135* were expressed under their respective duck promoters and used to generate PGCs expressing both genes separately. While the duck *RIG-I* promoter was previously described (22), we determined the activity of the duck *RNF135* promoter, which was examined by the generation of different deletion mutants tested in the NanoDLR^TM^ Assay System. Therefore, different deletion mutants were generated, including p1577, p1001, p349, and p197. Since we did not detect a core promoter activity of the duck *RNF135*, we used the full-length sequence of the duck *RNF135* for the generation of duck *RNF135*-expressing PGCs (Fig. 1a). The assembled expression vectors that were used to generate PGCs are presented in Fig. 1b.

**Figure 1.**
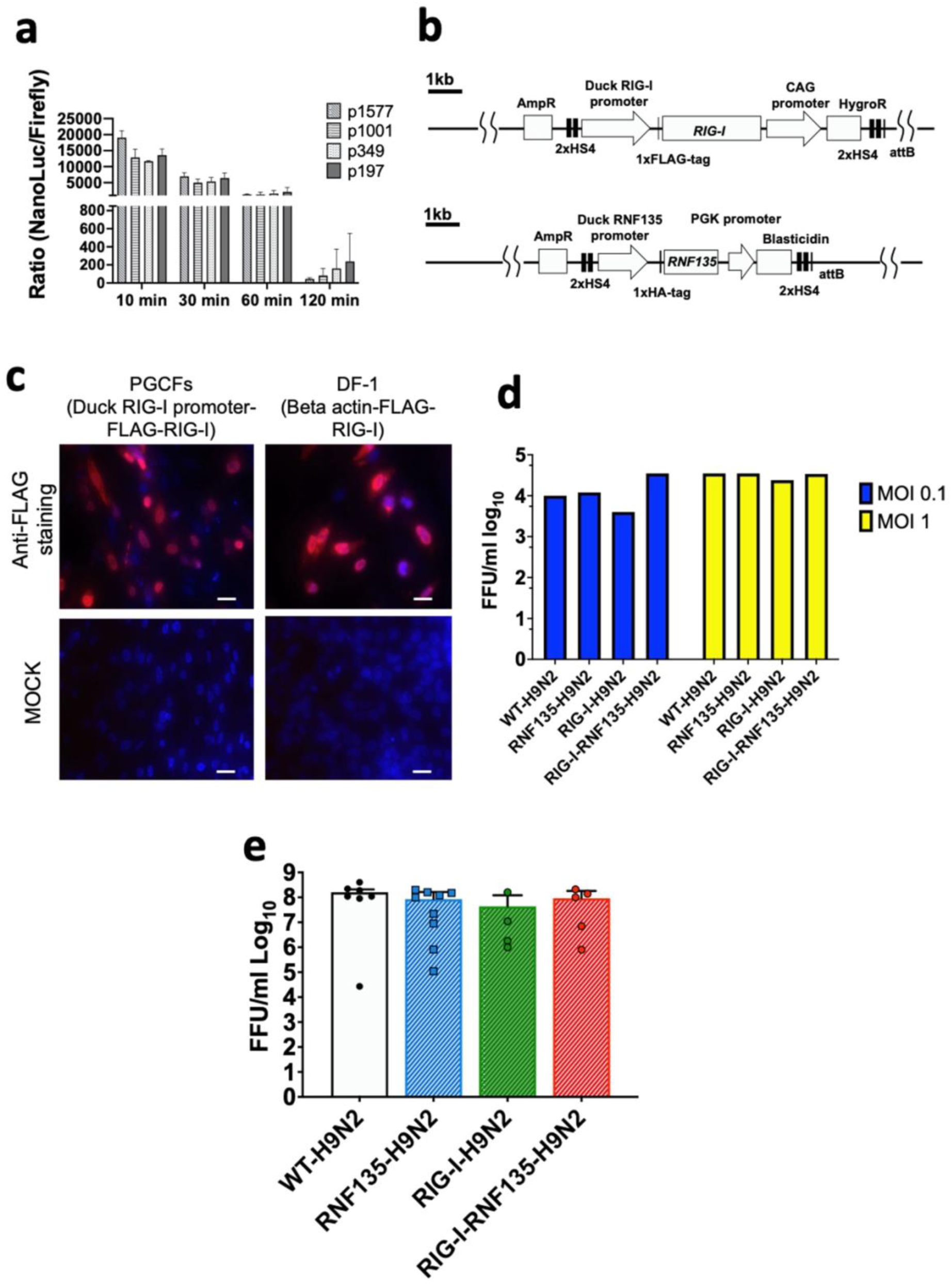
Generation of PGCs and susceptibility to infection using *in vitro* and *in vivo* systems. **a.** Promoter activity of the duck *RNF135* was examined by generating different deletion mutants tested in chicken DF-1 cells; the promoter activity was then evaluated by measuring the NanoLuc/Firefly ratio (n=3). **b.** Diagram of both constructs used to generate *RIG-I*-expressing chicken (upper diagram), using the previously identified duck *RIG-I* promoter (22) , and *RNF135*-expressing chickens (lower diagram), using the duck *RNF135* promoter, whose activity was identified in this study. **c.** Primordial germ cells (PGCs) that express the duck *RIG-I* under the duck *RIG-I* promoter were derived into PGC fibroblasts (PGCFs) and were infected later with avian influenza virus H9N2 to stimulate *RIG-I* expression upon influenza infection; the cells were infected for 18h with low pathogenic avian influenza virus H9N2 and subsequently stained for FLAG-Tag (red staining); MOCK control represents uninfected cells, and were not positive for anti-FLAG-Tag; DF-1 cells that express the FLAG-tagged duck *RIG-I* under the chicken beta-actin promoter were used as a positive control (red staining). **d.** Quantification of newly produced viral particles after infection of CEFs; CEFs were isolated from different transgenic embryos and experimentally infected with LPAIV H9N2 at two different multiplicities of infection (MOI 0.1 and 1); Supernatants were collected at 24hpi and titrated on MDCK cells; no significant differences were observed between the groups (*p*>0.05). **e.** Quantification of newly produced viral particles after infection of embryonated eggs. 10-day-old embryonated eggs were infected with LPAIV H9N2 at 10^3^ FFU/egg; allantois fluid was collected 24hpi and titrated on MDCK cells; no significant differences were observed between the groups (*p*>0.05). Error bars indicate the standard error of mean (SEM); Depending on the normal distribution of the data, multiple group comparison was done either with one-way ANOVA or Independent-Samples Kruskal-Wallis Test.

Due to the unavailability of commercial antibodies for detecting the duck *RIG-I*, we inserted a FLAG-Tag on the C-terminus to facilitate its detection using anti-FLAG antibodies. The activity of the duck *RIG-I* was examined by differentiating the *RIG-I*-expressing PGCs into PGC-derived fibroblasts (Fig. 1c, Supplementary Figure 1) that were infected with a low pathogenic avian influenza virus (LPAIV) H9N2 (Fig. 1c). Results showed that the infection led to activation of *RIG-I* as shown by FLAG-Tag staining. At the same time, the MOCK-infected cells remained negative for *RIG-I* expression (Fig. 1c).

### The expression of *RIG-I* did not limit the viral replication under *in vitro* and *in ovo* conditions

According to previously published data by Barber et al. (7), the overexpression of duck *RIG-I* in chicken DF-1 cells reduced the replication of H5N2 or H5N1 viruses when cells were infected at an MOI 1. Authors constitutively expressed *RIG-I* under the control of the CMV promoter using expression vector pcDNA 3.1. Therefore, we wanted to examine if the expression of *RIG-I* under the control of the duck RIG-I promoter can lead to a similar effect in limiting virus replication. To this end, we isolated chicken embryonic fibroblasts (CEFs) and produced embryonated eggs from the generated transgenic chickens and infected them with LPAIV H9N2. After virus infection, supernatants and allantoic fluid were collected from CEFs and embryonated eggs, respectively. We quantified the newly produced viral particles using a focus-forming assay. Surprisingly, transgene expression did not significantly affect the viral replication in both tested systems: CEFs (Fig. 1d, Supplementary Figure 2) and embryonated chicken eggs (Fig.1e, Supplementary Figure 3), which was the opposite in the previously published study (7). This highlights the importance of the chosen promoter, which may have affected the expression of *RIG-I* and influenced the innate immune response towards influenza virus.

### Phenotypic characterization of the genetically modified chickens

Upon the generation of *RIG-I-* and *RNF135*-expressing chicken lines, we wanted to ensure that the genetic modification did not negatively affect the growth of the generated transgenic birds. Therefore, we monitored their development by weekly measuring their body weights (Fig. 2a). Both generated chicken lines showed comparable growth to their WT siblings. The animals matured sexually, and no harmful phenotype was detected (Fig. 2a and 2b). The expression of both genes was examined via RT-PCR, revealing that both genes are expressed differentially in various tissues (Fig. 2c). We also detected comparable levels in expression levels with the duck (Fig. 2d)

**Figure 2.**
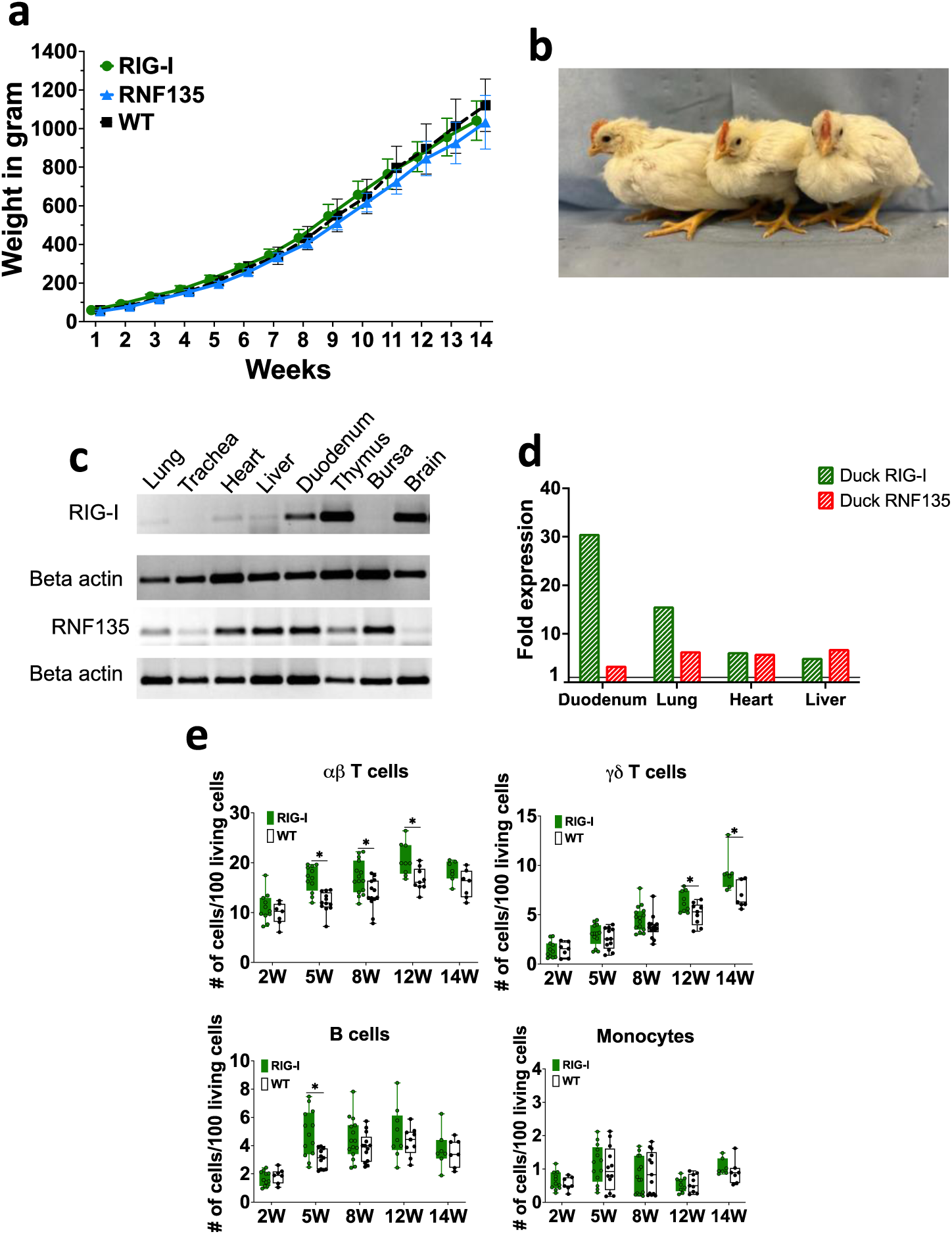
Generation and immunophenotypic characterization of *RIG-I* and *RNF135*-expressing chickens. **a.** Weekly weight monitoring of the generated heterozygous birds (*p*>0.05) (n=10). **b.** Representative picture of the generated heterozygous birds from left to right at four weeks of age: WT, *RNF135*-expressing chicken, and *RIG-I*-expressing chicken. **c.** RT-PCR of the transgenic expression of *RIG-I* and *RNF135* in different organs. **d.** Analysis of duck RIG-I and duck RNF135 expression in various tissues using reads per kilobase per million mapped reads (RPKM). **e.** Assessment of different immune cell populations in *RIG-I*-expressing chickens compared to their WT siblings. (^∗^) indicates statistical differences between groups tested simultaneously (*p*<0.05). Depending on the normal distribution of the data, two-group comparison was done with the Wilcoxon rank-sum test or two two-sample T-test

Previous studies show that mammalian *RIG-I* affects adaptive immunity, mainly T cells (17). Therefore, we sought to examine whether the re-expression of *RIG-I* in chickens will have similar effects. We analyzed the number of peripheral blood mononuclear cells (PBMCs) using flow cytometry to investigate the possible impact of expressing *RIG-I*, *RNF135,* or both on immune cell counts. No differences were observed in the number of immune cells between *RNF135*-expressing chickens and their WT siblings (Supplementary Figure 4). However, *RIG-I*-expressing chickens exhibited a significantly higher number of αβ and γδ T cells as well as B cells in comparison to their WT siblings (*p*<0.05) (Fig. 2e). This was not the case for monocyte counts, where no significant differences were observed (Fig. 2e). In addition, no significant differences were observed in *RIG-I*-expressing chickens compared to WT birds regarding the levels of IgM and IgY at 12 and 14 weeks of age (Supplementary Figure 5).

Further analysis of different T cell subpopulations indicated that *RIG-I*-expressing chickens had a significantly higher number of CD4+T cells, CD8α^neg^T cells and a significantly lower number of CD8α+^high^T cells (*p*<0.05) (Fig. 3a and 3b). T-cell activation was quantified after lectin activation using Concanavalin A (ConA). *RIG-I*-expressing chickens had a higher level of T cell activation in TCR1+/CD25+ T cells, which was not the case for TCR2,3+/CD25+ T cells (Fig. 3c).

**Figure 3.**
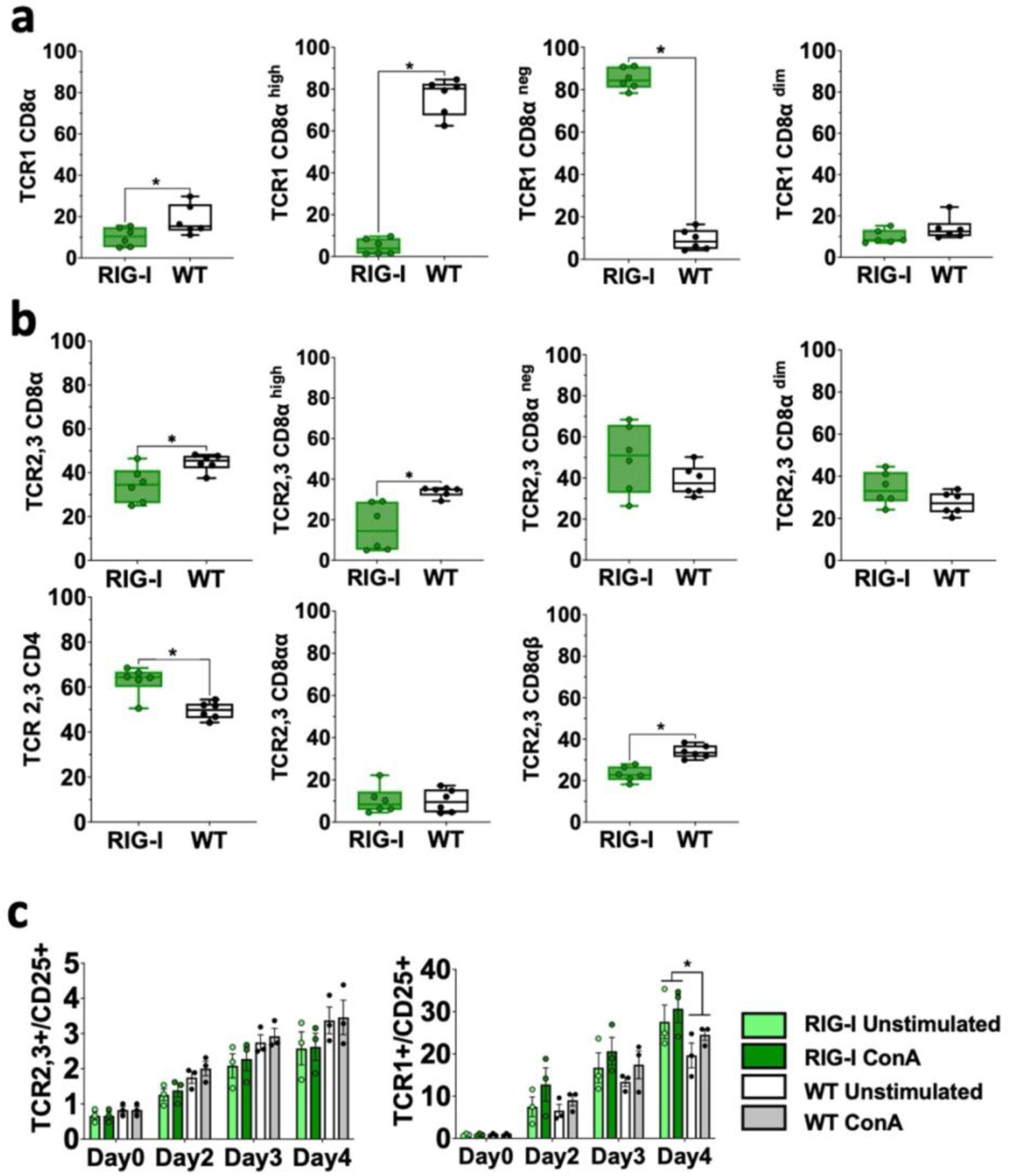
Assessment of different T cell subpopulations in *RIG-I*-expressing chickens compared to their WT siblings. **a.** PBMCS were analyzed for γδTCR1+/CD8α+ T cells at 12 weeks. B) PBMCS were analyzed for αβTCR2,3+T cells and CD4+ or CD8α+T cells at 12 weeks (*p*<0.05). **c.** Activation of isolated T cells from 12-week-old *RIG-I*-expressing chickens compared to their WT siblings (*p*<0.05); cells were sorted according to their TCR expression, stimulated with Concanavalin A, and quantified by flow cytometry at different time-points. The Y-axis depicts the number of positive cells per 100 viable TCR+ cells. Error bars indicate the standard error of mean (SEM); (^∗^) indicate statistical differences between groups tested simultaneously (*p*<0.05). Depending on the normal distribution of the data, two-group comparison was done with the Wilcoxon rank-sum test or two-sample T-test, while multiple group comparison was done either with one-way ANOVA or Independent-Samples Kruskal-Wallis Test

Heterozygous birds expressing each gene separately were crossed to obtain birds that simultaneously express both genes *RIG-I* and *RNF135*. The obtained *RIG-I-RNF135*-expressing chickens were monitored weekly for weight gain and closely investigated at eight and twelve weeks of age for possible alternate immune phenotypes similar to those observed in *RIG-I*-expressing birds. Unlike *RIG-I*-expressing chickens, no significant differences in T cells or B cell counts were detected in *RIG-I-RNF135*-expressing chickens in comparison to their WT siblings (Fig. 4d). Our analysis also comprised the investigated cell populations in *RIG-I*-expressing birds including αβ, γδ and B cells (Fig. 4a). Furthermore, we quantified CD4+, CD8+ T cells and corresponding subpopulations (Fig. 4b and Fig.4c). This was also the case for body weight gain, where no significant differences were observed in *RIG-I-RNF135*-expressing chickens in comparison to WT birds (Supplementary Figure 6).

**Figure 4.**
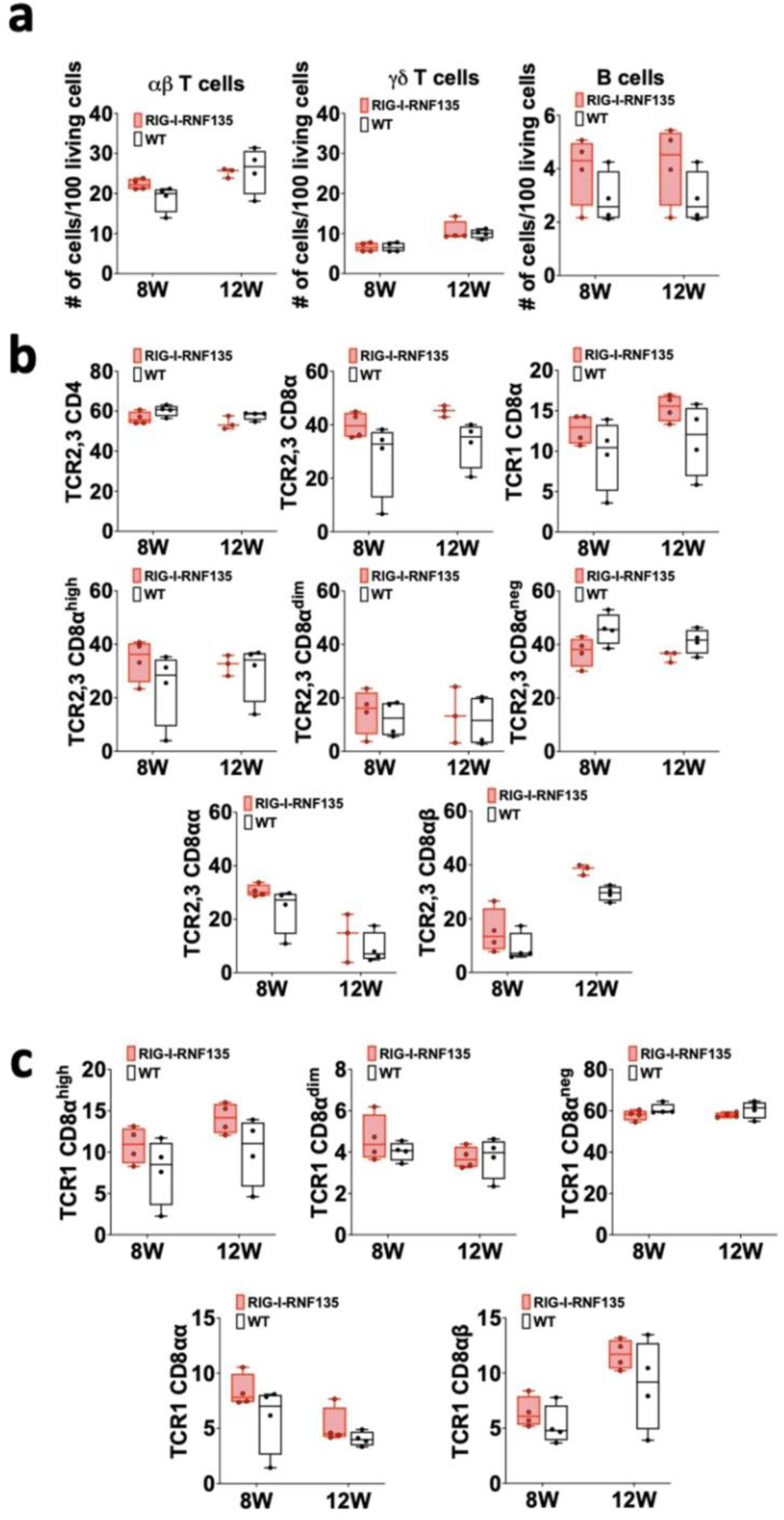
Simultaneous expression of *RIG-I* and *RNF135* in the chicken does not lead to differences in the adaptive immune phenotype compared to WT birds. Two-time points, eight and twelve weeks of age, were chosen based on the data obtained from *RIG-I*-expressing chickens to assess the immunophenotype of *RIG-I-RNF135*-expressing chickens. PBMCS were isolated and analyzed for B cells, αβTCR2,3+ or γδTCR1 + T cells (**a**) as well as for CD4+ (B) and CDα8+ T cells (**c**). No significant differences were detected between the analyzed groups (*p>*0.05). Depending on the normal distribution of the data, two-group comparison was done with the Wilcoxon rank-sum test or two-sample T-test

These data indicate that the *RIG-I*, an innate immune sensor, can influence adaptive immunity by causing shifts in T and B cell populations. In contrast, co-expression of *RNF135* with *RIG-I* seems to balance the adaptive immune cell populations, comparable to WT birds.

### *RNF135* is required for the potentiation of *RIG-I*’s activity

The role of *RNF135* as an essential ubiquitination factor that supports the antiviral activity of *RIG-I* was previously determined in mammalian cells (13). Later on, *RNF135* was found to be the obligatory ubiquitin for *RIG-I* (23), while so far, this was not known in birds. We investigated the effect of expressing *RNF135* on the antiviral activity of *RIG-I* in genetically modified chickens. We examined the susceptibility of the generated transgenic lines towards an H7N1 AIV, a direct precursor of a highly pathogenic avian influenza virus (HPAIV), known to cause severe acute respiratory disease in chickens (24). Birds were challenged with the virus and monitored for clinical signs, weight gain, viral replication, and lesion development. *RIG-I*-expressing chickens had the highest rate of morbidity, as indicated in Fig. 5a. The clinical symptoms started within the first two days of infection, where the morbidity rate reached a significant rate of 44% at one-day post-infection (dpi) (*p*<0.05) and increased to 67% at two dpi (*p*<0.05), which then decreased to 18% by three dpi (*p*<0.05). The onset of clinical disease was accompanied by a significant weight loss at two dpi that remained significantly low at six dpi (*p*<0.05) (Fig. 5c) and coincided with a pulmonary lesion score that increased from 0.8 at two dpi to 2.3 at six dpi (Fig. 6a and 6b). This was also confirmed by histological assessment of the caecum that revealed necrotic lesions associated with pronounced epithelial hyperplasia (Fig. 6b). *RIG-I-RNF135-* expressing chickens exhibited increasing clinical illness, from 13% of sick animals at one dpi to 50% at three dpi (*p*<0.05). Severe and prolonged clinical symptoms associated with mortality in this group lasted until six dpi.

**Figure 5.**
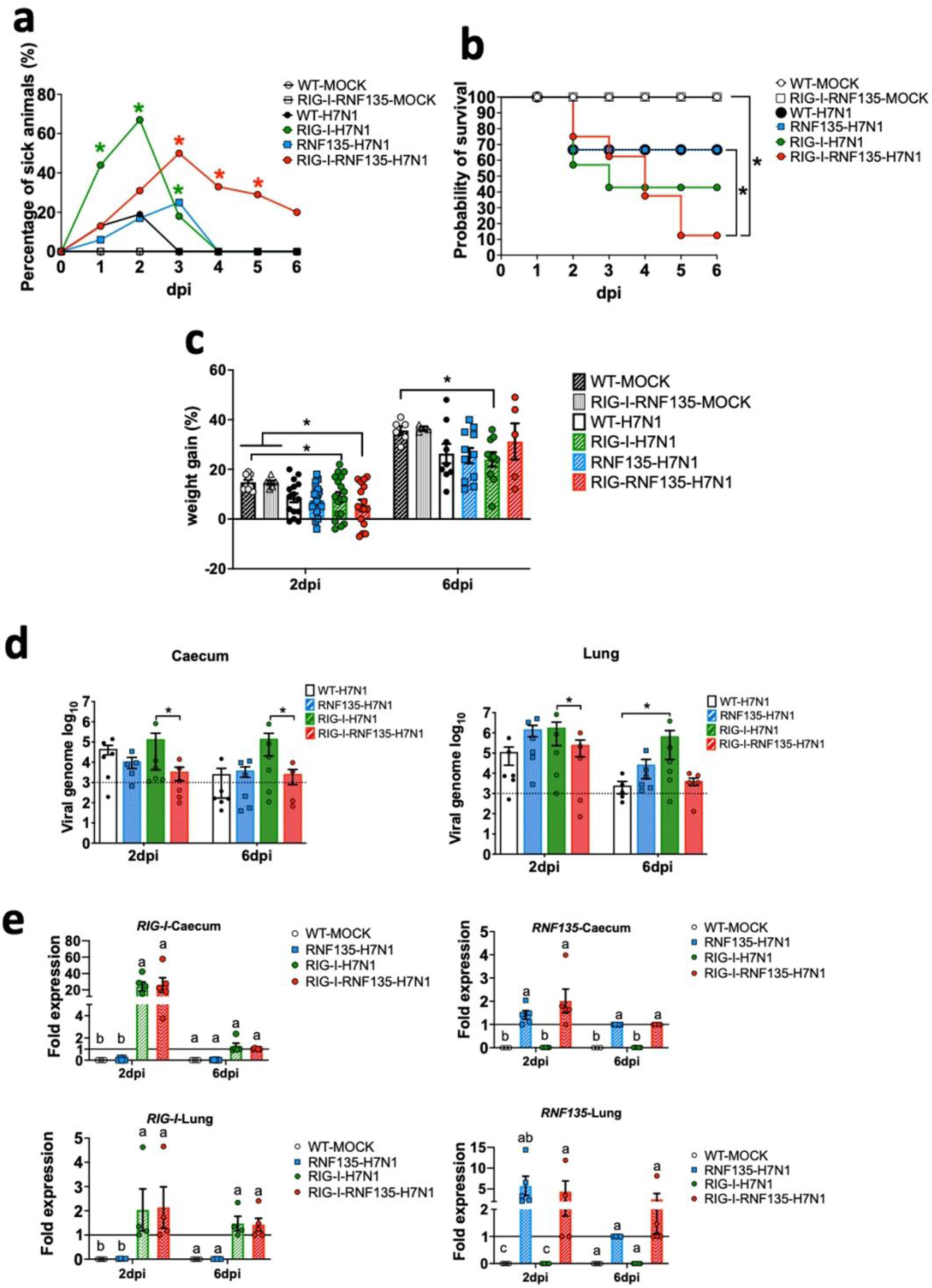
H7N1-challenge experiment reveals the susceptibility of *RIG-I*-expressing chickens and the role of *RNF135* in effective *RIG-I* antiviral response. The generated transgenic chickens were challenged with H7N1 and assessed at two days post-infection (dpi) and six dpi for different parameters. **a.** Percentage of animals presenting clinical symptoms. **b.** Probability of survival in the challenged groups. **c.** Weight gain in challenged groups compared to WT-and *RIG-I-RNF135*-MOCK-controls (*p*<0.05). **d.** Viral replication rate in two main target organs, caecum, and lung (*p*<0.05); the horizontal line indicates the detection threshold of the PCR based on the signal obtained from the uninfected controls, which is log_10_(10^3^). **e.** Expression levels of transgenes in the caecum and the lung upon H7N1-challenge. Error bars indicate standard error of mean (SEM); (^∗^) or different letters indicate statistical differences between groups tested simultaneously (*p*<0.05). Depending on the normal distribution of the data, multiple group comparison was done either with one-way ANOVA or Independent-Samples Kruskal-Wallis Test

**Figure 6.**
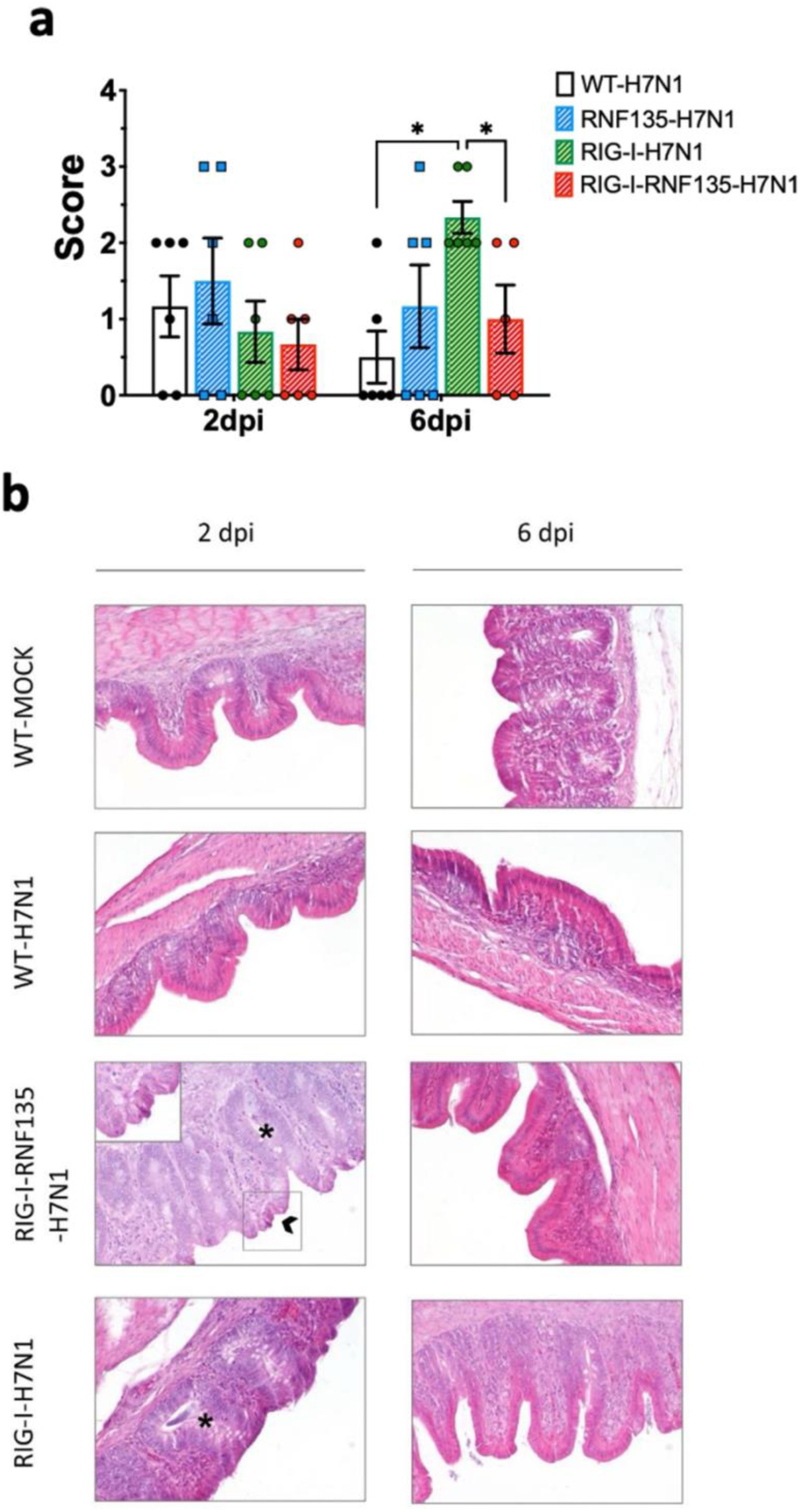
Developed lesions in H7N1-challenged birds. **a.** Macroscopical lesion score of the lungs showing a significant increase of lesions in *RIG-I*-expressing birds at six dpi (*p*<0.05) **b.** Histology of the caecum showing the typical structure of the epithelium in the control groups. *RIG-I-RNF135* challenged birds showed epithelial hyperplasia with necrosis that diminished by 6 dpi. *RIG-I*-expressing chickens did not show necrotic lesions but pronounced epithelial hyperplasia that lessened by six dpi. Asterix indicates epithelial hyperplasia with the mitotic figures, while the arrow indicates necrotic epithelial cells; 200x. Error bars indicate standard error of mean (SEM); (^∗^) indicate statistical differences between groups tested simultaneously (*p*<0.05). Depending on the normal distribution of the data, multiple group comparison was done either with one-way ANOVA or Independent-Samples Kruskal-Wallis Test

The expression of *RNF135* alone resulted in a mortality rate comparable to WT birds. In contrast, the clinical symptoms in *RNF135*-expressing chickens lasted till three dpi with a total of 25% sick animals (Fig. 5b). This was not the case for WT chickens that exhibited clinical symptoms only for the first two days after infection and had a morbidity rate of 13% and 19% at one and two dpi, respectively (Fig. 5b).

The quantification of the virus genome copies via qRT-PCR showed that the expression of *RIG-I* together with *RNF135* significantly reduced the amount of viral genome copies in the caecum compared to *RIG-I* expressing chickens at two and six dpi (*p*<0.05) (Fig. 5d). In the lung, challenged *RIG-I-RNF135*-expressing chickens displayed a virus replication rate similar to WT birds and a significantly lower replication rate than *RIG-I*-expressing chickens at two dpi (*p*<0.05). Conversely, *RIG-I*-expressing chickens had the highest viral replication rate in the lung and the caecum among all investigated groups, with a significant difference compared to *RIG-I-RNF135*-expressing chickens (*p*<0.05) (Fig. 5d). The amount of viral nucleic acid remained at a high level at six dpi, indicating the lack of viral clearance in these birds, which was not the case for the other challenged chicken lines (Fig. 5d).

These results indicated that the reinstatement of *RIG-I* or *RIG-I*-*RNF135*-expressing chickens had no effect on viral replication compared to WT birds. The different outcome in viral replication in *RIG-I-RNF135*-expressing chickens, compared to *RIG-I*-expressing chickens, confirms the role of *RNF135* as a ubiquitin ligase at the early stage of infection. Also, it demonstrates that the chicken *TRIM25* does not overtake the role of *RNF135* in *RIG-I*-expressing chickens.

### The expression of *RIG-I* and *RNF135* in H7N1-challenged birds coincided with the acute phase of infection

We aimed to quantify possible changes in the expression levels of both transgenes in the challenged animals after the H7N1 infection. We quantified *RIG-I* and *RNF135* expressions via qRT-PCR in the lung and the caecum (Fig. 5e). The caecal expression of *RIG-I* in virus-infected *RIG-I-RNF135* chickens was increased by ∼26 fold. At the same time, infected *RIG-I* chickens exhibited an increase of ∼24-fold compared to *RIG-I-RNF135* MOCK controls. Infected *RIG-* and *RIG-RNF135-*expressing chickens had a comparable *RIG-I* expression to *RIG-RNF135*-MOCK controls with no significant upregulation (Fig. 5e). Overall, we observed lower expression of *RNF135* in the caecum and lungs compared to *RIG-I-RNF135*-MOCK controls. The expression increased by 1.4 fold in *RNF135*-infected birds and ∼2 fold in *RIG-I-RNF135*-infected chickens. In the lung, the *RNF135* expression at two dpi was upregulated by ∼6 fold and ∼4 fold in infected *RNF135* and *RIG-I-RNF135*-expressing chickens, respectively. At six dpi, the *RNF135* expression was at a comparable level with the *RIG-I-RNF135* MOCK controls (Fig. 5e). These data indicate that the infection led to a quick upregulation of *RIG-I* at the acute phase of infection, which quickly decreased afterward.

### Differential regulation of innate immune genes in naïve as well as in H7N1-challenged *RIG-I*, *RNF135,* and *RIG-I-RNF135*-expressing chickens

We speculated that the observed acute inflammatory reaction could be due to a differential regulation of inflammatory genes related to *RIG-I* signaling. Therefore, we sought to quantify the differentially expressed genes (DEGs) between transgenic and WT birds. We used the Fluidigm qRT-PCR array to identify changes in the expression of selected innate immune genes in both naïve and H7N1-challenged birds (25). The results revealed that in the absence of infection, *RIG-I-RNF135*-expressing chickens already had a significantly higher expression of ISGs in the caecum, including *ISG12-2* and *OASL,* as well as the transcription factor Early Growth Response *1 (EGR1)* (Fig. 7a, Supplementary Figure 7). In contrast, others, including *EGR1*, interleukin 4 induced 1 (*IL4I1)*, Interleukin-1 beta (*Il1β),* and Interleukin 8 (*IL8),* were significantly upregulated in the spleen (Fig. 7a and Supplementary Figure 7).

**Figure 7.**
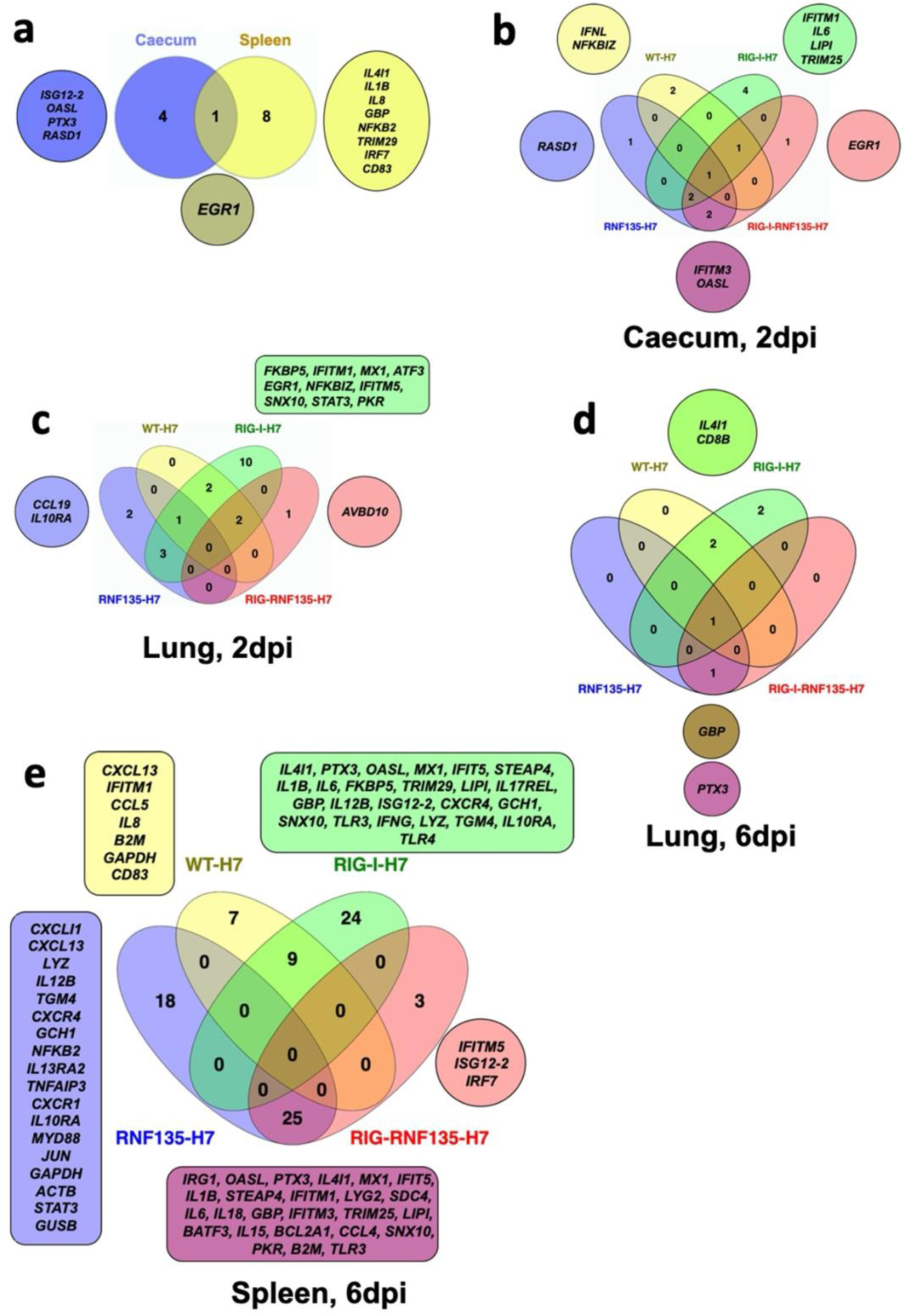
Infection of transgenic chickens with H7N1 leads to upregulated inflammatory genes. Venn diagram of fluidigm qRT-PCR array of naive and challenged birds with H7N1 showing significantly upregulated genes in various organs. Significant DEGs were identified by comparing the relative expression values for every chicken line to the WT-MOCK individually per timepoint, with a significance level set at *p* < 0.05; fold change >1 (n ≥ 5-time point). **a.** Upregulated genes in *RIG-I-RNF135*-expressing chickens without infection **b.** Upregulated genes in the caecum between H7N1-challenged groups at 2dpi **c.** Upregulated genes in the lung between H7N1-challenged groups at 2dpi **d.** Upregulated genes in the lung between H7N1-challenged groups at 6dpi **e.** Upregulated genes in the spleen between H7N1-challenged groups at 6dpi

The comparison of challenged groups with non-infected controls (WT-MOCK) indicated that *RIG-I* chickens expressed several inflammatory genes in the caecum and lungs that were not expressed in the other groups. This included *IFITM1*, *IL6,* and *LIPI* (Fig. 7b, Supplementary Figure 7c and 7d). At two dpi, we observed the upregulation of genes involved in the JAK-STAT signaling pathway, including *IL-6* and *STAT3,* which was not the case at 6 dpi (Fig. 7b, 7c, and 7d). The significant upregulation of inflammatory and immune regulatory genes in the *RIG-I*-expressing chickens persisted until six dpi, where over 20 genes were exclusively upregulated in the spleen (Fig. 7e and Supplementary Figure 7e). This was not the case for *RIG-I-RNF135*-expressing chickens and WT chickens with an exclusive expression of three and seven genes, respectively (Fig. 7e and Supplementary Figure 7e). In addition, the analysis of expressed genes in the spleen revealed that only *RIG-I-RNF135* chickens expressed *IRF7* and *TRIM25*, in contrast to *RIG-I*-expressing chickens (Fig. 7e and Supplementary Figure 7e).

The functional enrichment analysis of the genes involved in the biological processes of transgenic chickens indicated that the regulated genes in *RIG-I*-expressing chickens were highly involved in metabolic activities. In contrast, those in *RIG-I-RNF135*-expressing birds were primarily involved in regulatory mechanisms (Supplementary Figure 8). Overall, the obtained data confirm that *RIG-I*-expressing chickens exhibited a significant increase of inflammatory cytokines that were not observed in other challenged birds, which may explain the acute inflammatory reaction in these birds.

### High expression of *IFN-γ* is associated with significant virus replication in *RIG-I*-expressing birds

Since we observed a significant reduction of viral genome copies in the *RIG-I-RNF135*-expressing chickens compared to *RIG-I*-expressing birds, we compared the DEGs between H7N1-infected groups. This helped us determine possible factors behind the increased H7N1 replication in *RIG-I*-expressing birds. Interestingly, these birds lacked interferon upregulation compared to infected WT birds or *RIG-I-RNF135*-expressing chickens (Fig. 8a). We also found that the viral infection led to a significant increase in the expression of *IFN-γ* at six dpi when compared to H7N1-infected *RIG-I-RNF135*-expressing chickens (14*-*fold change increase) and WT birds (9-fold change increase) (Fig. 8b and 8d) (*p*<0.05). In contrast, *RIG-I-RNF135*-expressing birds quickly exhibited a significant increase of *IFN-α* expression, already at two dpi, in comparison to *RIG-I*-expressing birds (12-fold change) as well as to WT birds (17-fold change) (Fig. 8c). We concluded that the co-expression of *RNF135* and *RIG-I* in chickens leads to a significant increase of *IFN-α* expression, which was not observed when *RIG-I* was expressed solely. In addition, *RIG-I*-expression caused an upregulation of *IFN-γ* (Fig. 8b and 8d) that was accompanied by a significant increase in virus genome copies.

**Figure 8.**
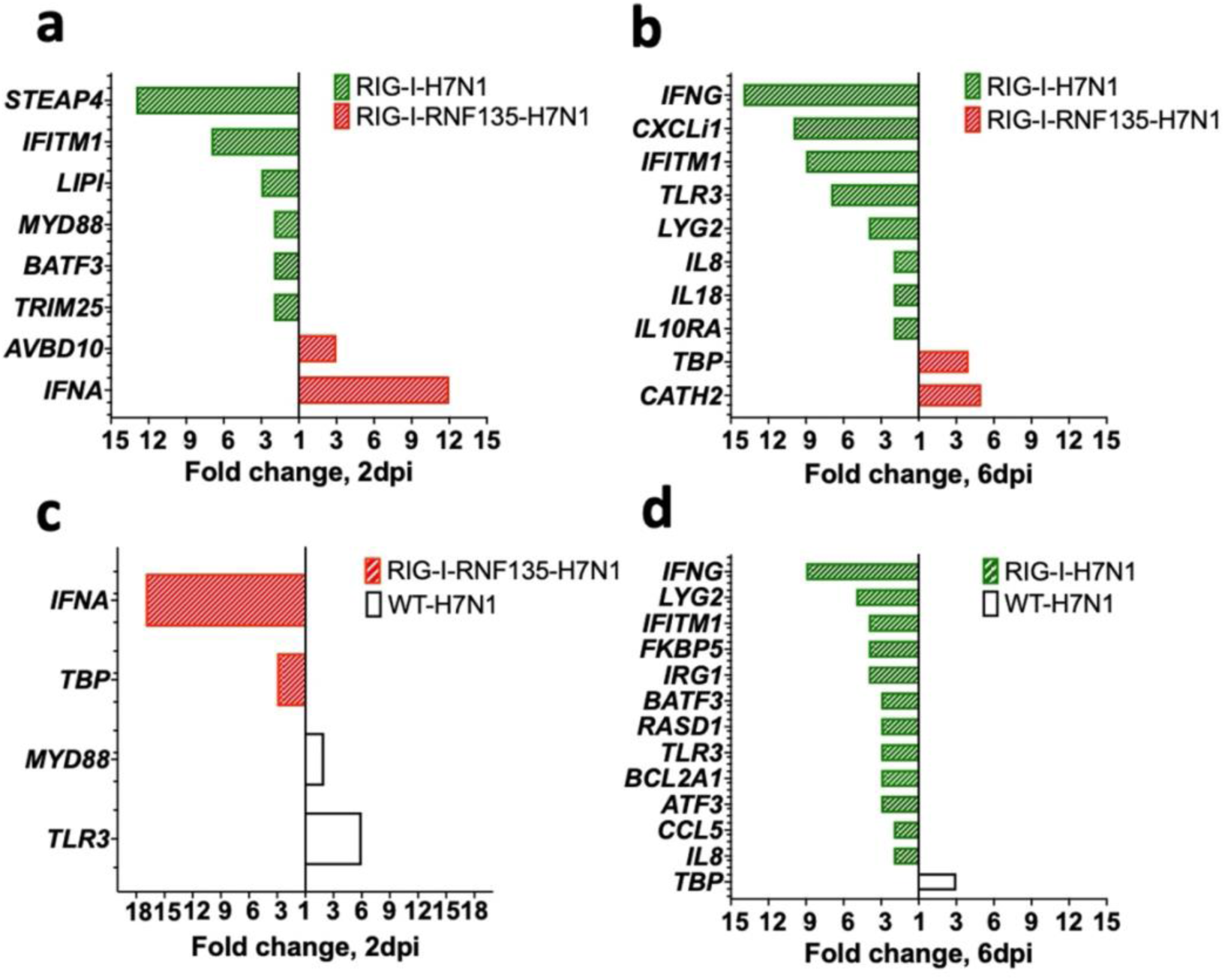
Gene expression in the lung upon H7N1-infection. Significant DEGs were identified by using Fluidigm qPCR array and comparison of the relative expression values between H7N1-challenged groups. **a.** Significantly upregulated genes after comparison of infected *RIG-I*-with *RIG-I-RNF135*-expressing chickens at two dpi. **b.** Significantly upregulated genes after comparison of infected *RIG-I*-with *RIG-I-RNF135*-expressing chickens at 6dpi. **c.** Significantly upregulated genes after comparison of infected *RIG-I*-with *RIG-I-RNF135-*expressing chickens at two dpi. **d.** Significantly upregulated genes after comparison of infected *RIG-I*-expressing with infected WT-chickens at six dpi. Significance levels were set at *p* < 0.05; fold change >1 (n ≥ 5-time point)

### The deleterious inflammatory response in *RIG-I* and *RIG-I-RNF135*-expressing chickens depends on the virulence of the influenza subtype

The infection with H7N1 revealed a unique phenotype that manifested in acute inflammation and increased mortality in *RIG-I* and *RIG-I-RNF135*-expressing chickens. Hence, we were interested in investigating the effect of reinstating *RIG-I* and *RNF135* in the chicken genome on the outcome of infection with two additional virus strains of high or low virulence. We conducted two additional *in vivo* infection studies using two viruses: an H3N1 (A/chicken/Belgium/460/201) and an H9N2 (A/chicken/Saudi Arabia/CP7/1998). While both subtypes are classified as low pathogenic AIVs due to their monobasic hemagglutinin cleavage sites, H3N1 has been described as a highly virulent strain causing severe clinical infection and mortality in adult layers (26). Displaying a distinct tropism for the hen’s oviduct, H3N1 causes salpingitis and peritonitis, which are associated with a severe drop in egg production. In contrast, the H9N2 virus is effectively low virulent and does not cause any detectable symptoms in experimentally infected chickens(27). The H9N2-and H3N1-infection experiments of *RIG-I* and *RIG-I-RNF135*-expressing chickens revealed major differences in the outcome of infection between the two viruses. While H9N2 infection did not cause any clinical disease in the *RIG-I* or *RIG-I-RNF135*-expressing chickens (Supplementary Figure 9), infection with H3N1 led to marked clinical/pathological disease signs and disease aggravation, similar to those observed in the H7N1 infection experiment. The infection with H3N1 led to early clinical disease and mortality onset that were more pronounced in *RIG-I*-expressing birds (Fig 9a, 9b and Supplementary Figure 10). However, we did not detect differences in viral RNA loads in tracheal swabs between the H3N1-infected groups except for 7dpi, where *RIG-I*-expressing birds exhibited significantly higher loads compared to WT birds (Fig. 9a). *RIG-I*-expressing chickens also manifested a significantly increased expression of *IL-1β*, *IL6*, *IFN-α* and *IFN-γ* compared to the other infected birds (Fig. 9c). The assessment of histological lesions of the reproductive system indicated that the infection with H3N1 led to pronounced atrophy in *RIG-I* and *RIG-I-RNF135*-expressing birds in comparison to WT birds (Fig. 9d), accompanied by fibrinous peritonitis, salpingitis and vasculitis (Fig. 9e). These data confirm the observations made with H7N1 regarding the harmful inflammation. They indicate that the deleterious clinical disease caused by *RIG-I* reinstatement depends on the degree of viral virulence.

**Figure 9.**
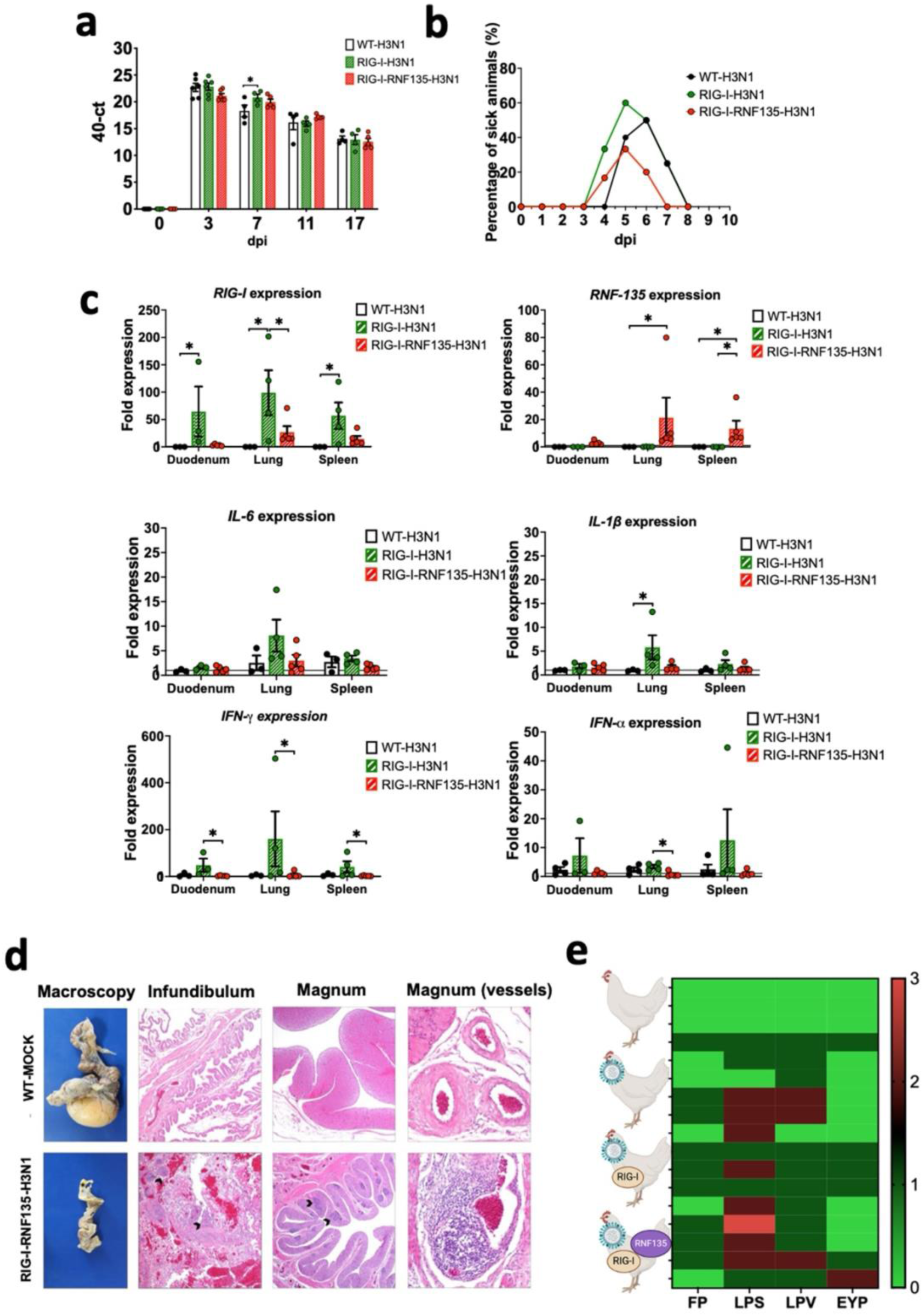
H3N1-challenge experiment confirms the exacerbated disease phenotype of *RIG-I* and *RIG-I-RNF135* chickens infected with virulent avian influenza viruses. The generated transgenic chickens were challenged at 28 weeks of age with H3N1 and assessed for different parameters. **a.** Viral shedding based on tracheal swabbing and viral RNA load analysis. b. Probability of survival in the challenged groups. **c.** Expression of *RIG-I*, *RNF135*, and influenza-regulated genes in the duodenum, lung, and spleen. **d.** Histological assessment of the reproductive tract; WT-MOCK: normal macroscopical and histological appearance of the salpinx; RNF-RIG-I-H3N1: severe atrophy with mild fibrinous peritonitis in the infundibulum (asterisk), severe lymphoplasmacytic salpingitis (arrowheads) (40x), the magnum (20x), and vasculitis in the magnum vessels (200x). **e.** Scoring of lesions in the reproductive tract in all challenged groups, starting with the upper row WT-MOCK, WT-H3N1, *RIG-I*-H3N1, and *RIG-I-RNF135*-H3N1; FP: fibrinous peritonitis, LPS: lymphoplasmacytic salpingitis, LPV: lymphoplasmacytic vasculitis, EYP: egg yolk peritonitis. Error bars indicate standard error of mean (SEM); (^∗^) indicate statistical differences between groups tested simultaneously (*p*<0.05). Depending on the normal distribution of the data, multiple group comparison was done either with one-way ANOVA or Independent-Samples Kruskal-Wallis Test

## Discussion

The constant arms race between pathogens and the host affects different aspects of the immune system, including innate sensors. The reason for the loss of *RIG-I* in Galliformes, which correlated with the disappearance of the ubiquitin ligase *RNF135*, remained a mystery (20), especially given the established contributing role of *RIG-I* in the resilience towards influenza in ducks and other species (7, 28). It was previously speculated that the gradual loss of *RIG-I* and *RNF135* in the chicken was possibly caused by the pathogen’s resistance to the innate sensor or the disappearance of some relevant pathogens (20). At the same time, preservation of the antiviral competence accompanied the loss of *RIG-I* and *RNF135* in chickens via the evolutionary selection of *MDA5* and *LGP2* (29). The beneficial effect of the duck *RIG-I-*overexpression in chicken DF-1 cells was demonstrated by the limited replication of AIV in these transgene-expressing cells (7). So far, no chickens expressing the duck *RIG-I* have been generated to investigate their antiviral response *in vivo*. In this study, we produced genetically modified chickens that express *RIG-I* with or without its ubiquitin ligase *RNF135* to examine the physiological role of both genes and their function during AIV infection. Both genes were cloned from the duck representing the most studied avian influenza reservoir with evolutionary conserved *RIG-I* and *RNF135* (6, 20, 30). The expression of both genes was controlled under their respective duck promoters since the duck *RIG-I* was previously tested in chicken cells(22). This strategy allows the strict control of gene expression that can be induced only upon infection, which prevents undesirable production of inflammatory cytokines that may cause autoimmune diseases (22). While the duck *RIG-I* promoter was previously identified (22), we described the activity of the duck *RNF135* promoter using chicken cells.

Without infection, *RIG-I* expression did not cause any harmful phenotype, but led to different adaptive immune cell counts compared to WT siblings. This suggests the role of the *RIG-I* in priming T cell immunity in the chicken, which can be similar to mice that exhibited a lack of T cell immunity associated with deficiencies in migratory dendritic cell activation, viral antigen presentation and CD8+ and CD4+ T cell priming (17) (14). In the case of *RIG-I*-*RNF135*-co-expression, the balanced adaptive immune phenotype implies a possible role of *RNF135* in modulating the T cell immune response in birds, which was previously described for Th1 response and cytotoxic T cells in mice (31). Moreover, naïve *RIG-I-RNF135*-expressing chickens had a significantly higher expression of interleukin 4 inducible gene 1 (*IL4I1*) as well as protein nuclear translocation 7 *(IRF7*) and tripartite motif-containing protein 29 *(TRIM29)*. This may explain the differential adaptive immune phenotype between *RIG-I* and *RIG-I-RNF135*-expressing chickens. Previous studies indicated the involvement of *IL4I1* in the signaling to T and B lymphocytes (32) and the effective role of translocation factors such as *IRF7* in chicken cells transfected with the duck *RIG-I* (22).

The lack of an antiviral effect after *in ovo* and *in vitro* experimental infections with LPAIV H9N2 was discordant with the previously described antiviral effect of duck *RIG-I* in chicken DF-1 cells (7). Barber et al, observed low virus replication upon the overexpression of *RIG-I* under the control of the CMV promoter. The difference between our results and previously published data can be related to the chosen promoter used by Barber et al, which may lead to a higher upregulation of *RIG-I* and, consequently, a strong production of interferon-stimulated genes. Additional factors can stand behind the differences between the previously published results by Barber et al(7), and our data, including influenza subtypes and the type of cells.

H7N1-challenge experiments of the generated transgenic birds revealed that *RIG-I*-expressing chickens had an early upregulation of proinflammatory cytokines such as *IL-6*, known to support *IL1β* in suppressing regulatory T cells, which can lead to an uncontrolled increase in the number of CD4+ T cells (33). Furthermore, the expression of *RIG-I* without *RNF135* was not beneficial in limiting H7N1 replication since the *RIG-I*-expressing chickens had the highest viral genome copies among all challenged groups. The requirement of *RNF135* for an *RIG-I*-mediated antiviral effect in *RIG-I-RNF135*-expressing chickens was also described in mammalian cells (13). In addition, we found that the H7N1 infection caused a significant upregulation of *RIG-I* in the caecum compared to the lungs. This can be because *RIG-I* expression rapidly occurs upon stimulation and may reach a peak by three hours post-infection (34). Similar observations were described by Cornelissen et al. that detected a significant upregulation of *RIG-I* expression in the lungs of ducks at 8h post H7N1 infection that significantly decreased at two dpi (35). *RIG-I*-expressing birds infected with H7N1 exhibited an acute inflammatory response and weight loss at two dpi, lasting till six dpi. Pang et al, reported that influenza virus could hijack the inflammatory reaction associated with RIG-I signaling for its replicative advantage, particularly in the respiratory tract (15). H7N1-infected *RIG-I*-expressing birds showed several regulated immune genes compared to other infected groups, including upregulation of the suppressor of cytokine signaling 1 (*SOCS1*) in the caecum. This may explain the significant increase in viral genome copies and the observed T cell phenotype since *SOCS1* is a potent inhibitor of JAK/STAT signaling (36) and is involved in several mechanisms that regulate the T cell maturation, differentiation, and function (37). Furthermore, the assessment of the immunophenotype of *RNF135*-expressing chickens revealed differential upregulation of various genes, which suggests that this gene, on its own, may function independently from *RIG-I* (38).

Interestingly, we detected significant upregulation of *IFN-γ* in the lungs of infected *RIG-I*-expressing chickens compared to infected *RIG-I-RNF135-* and WT-chickens, which can be responsible for lung-mediated injury and acute death caused by respiratory distress. Similarly, H3N1 infection caused a relative upregulation of *IFN-γ* in the spleen and the duodenum of *RIG-I*-expressing chickens. The role of *IFN-γ* was previously described in *IFN-γ* KO mice that were less susceptible to lung inflammation and pathology upon influenza infection (39). The lack of an antiviral effect in *RIG-I*-expressing chickens may also indicate that chicken *TRIM25* is dispensable for *RIG-I* efficient ubiquitination. Similar findings were described using human lung adenocarcinoma epithelial cells, where *TRIM25* did not participate in the endogenous *RIG-I*-dependent antiviral responses (21). However, other studies done in mammalian models indicated that the deletion of *TRIM25* increases the susceptibility to influenza infection (40), supporting the evidence that *TRIM25* may bind directly to the viral RNA, thereby contributing to the restriction of influenza virus infection (41). The exacerbated inflammatory reaction in *RIG-I*-expressing chickens could be related to the absence of *RNF135*, which suggests a stabilizing role of *RNF135* comparable to *TRIM25* (42), though this requires further investigation. The increased mortality rates in *RIG-I-* and *RIG-I-RNF135*-expressing chickens may be explained by immunopathology in both chicken lines, despite the differences in viral replication. The additional *in vivo* experimental challenge with the highly virulent H3N1 virus confirmed the deleterious immunophenotype observed for the H7N1 virus, as demonstrated by the upregulation of various inflammatory genes, including *IFN-γ, IFN-α, IL1β, and IL6*. The absence of a similar phenotype after infection with the mildly virulent H9N2 virus revealed that disease exacerbation in *RIG-I*- and *RIG-I-RNF135*-expressing chickens requires a certain degree of viral virulence.

Our data suggest that the evolutionary loss of *RIG-I* in chickens and other galliform birds was advantageous in coping with viral infections caused by AIV or other avian RNA viruses. This subsequently helped decrease the acute inflammation and the damage to the host. A comparable hypothesis was made in the case of pangolins that lost the *MDA5* as an evolutionary mechanism to cope with coronavirus-induced inflammation (43). The acute inflammation seen in the generated chickens may be linked to the duplicated function in RNA sensing due to the positive selection of *MDA5* (20), which could exacerbate the inflammatory response and requires further investigation. Above that, we propose that the antiviral role of *RIG-I* in ducks (7) is not exclusively related to this gene. Still, it may involve the ubiquitination factor *RNF135* and possibly other unknown regulatory factors that support *RIG-I* signaling and reduce the repercussions of acute inflammation by negatively regulating *RIG-I* signaling (44). We provided novel information regarding the outcome of the re-introduction of *RIG-I* and its ubiquitination factor *RNF135* in the chicken genome. Future work should focus on identifying factors that can help reduce the acute inflammatory reaction in *RIG-I-RNF135*-expressing chickens while maintaining potent antiviral activity, which can lead to the generation of avian influenza virus-resistant chickens.

## Materials and Methods

### Cloning of the duck *RIG-I* and the duck *RIG-I* promoter

Total RNA was isolated from the spleen of the mallard duck (*Anas platyrhynchos*) using ReliaPrep^TM^ RNA Tissue Miniprep System (Promega, USA), followed by cDNA synthesis using GoScript™ Reverse Transcription System (Promega, USA) according to the manufacturer’s instructions. The duck *RIG-I* was amplified using Q5® High-Fidelity DNA Polymerase (New England Biolabs, USA) using the primers 562_RIG-I_for (5’-ATGACGGCGGACGAGAAGCGGAGC-3’) and 563_RIG-I_rev (5’-CTAAAATGGTGGGTACAAGTTGGAC-3’) that were previously described (45). The PCR thermal conditions were as follows: 98 °C 30 sec, followed by 35 cycles of 98 °C 10 sec, 67 °C 20 sec, 72 °C 1:30 min, and a final extension step of 72 °C 2 min.

The entire length of the duck *RIG-I* promoter was amplified using primers 706_RIG-I_For (5’-AGCTGATGACCTGCAAAAAGTT-3’) and 661_RIG-I_Rev (5’-GGCTGGGCTCTGCCGGCCG-3’), which were described elsewhere (22). This resulted in an amplicon of 2017 bp that was fully sequenced and aligned with the duck genome (*Anas platyrhynchos*, NC_040075.1). The PCR was conducted following the following thermal conditions 98 °C 30 sec, followed by 35 cycles of 98 °C 10 sec, 70 °C 20 sec, 72 °C 1:30 min, and a final extension step of 72 °C 2 min.

### Cloning of the duck *RNF135*

The genomic region containing duck *RNF135* was obtained from the GenBank contig PEDO01000017.1 and corrected based on multiple publicly available duck RNAseq data. Although the *RNF135* sequence was correctly predicted in many birds, we detected a partially incomplete annotated sequence of the duck *RNF135*, specifically the missing 5‘ part that contains the RING domain (previous accession number XM_013092775), while the predicted version of the *RNF135* was made available (XM_027471415.2). The full-length sequence of the duck *RNF135* was synthesized after codon optimization using the IDT online tool (IDT™, USA) (Supplementary Figure 11). The obtained *RNF135* sequence was subsequently cloned into the *RNF135*-expression vector driven by the duck *RNF135* promoter (Fig. 1b).

### Identification of the duck *RNF135* promoter via Nano-Glo^®^ Dual-Luciferase^®^ Reporter Assay

The putative duck *RNF135* promoter was obtained by amplifying 1577 bp 5’ of the ATG start codon from duck gDNA cloned into pGEM vector (Promega, USA) and analyzed by Sanger sequencing. The PCR was done using Q5® High-Fidelity DNA Polymerase with primers: 707_RNF_Prom_For (5’-GA GCA GAG CCA GGC AGC TAT A-3’), 708 (5’-GGT CCT GCT CGG GGC GGA GC-3’) resulting in an amplicon of 1557 bp. The PCR thermal conditions were conducted using Q5® High-Fidelity DNA Polymerase (New England Biolabs, USA) at two step-PCR 98 °C 30 sec, 98 °C 10 sec, 72 °C 60 sec and a final elongation step of 72 °C 2 min.

The promoter activity of duck *RNF135* was assessed by measuring promoter-driven NanoLuc^TM^ luciferase activity normalized to the luminescence of Firefly luciferase. To this end, a total of 50.000 chicken DF-1 cells were seeded in 24 well plates and were co-transfected 24h later with a vector plasmid containing the deletion mutant and a second plasmid for the expression of Firefly under the PGK promoter (Promega, USA). 24h after transfection, cells were washed with PBS, trypsinized, and resuspended in 250µl culture medium. Firefly signal was detected by mixing 80µl of the cell suspension with the same amount of ONE-Glo^TM^ EX Reagent, prepared by combining ONE-Glo™ EX Luciferase Assay Buffer with ONE-Glo™ EX Luciferase Assay Substrate in 1:1 ratio (Promega, USA). After measuring the signal of the Firefly luciferase in the GloMax® 20/20 Luminometer (Promega, USA), 80 µl NanoDLR™ Stop & Glo® reagent, prepared by adding NanoDLR™ Stop & Glo^®^ Substrate 1:100 into NanoDLR™ Stop & Glo® Buffer (Promega, USA), were added to the samples. These were incubated for 10, 30, 60, and 120 min and measured again in the GloMax^®^ 20/20 Luminometer (Promega, USA) to detect the NanoLuc^TM^ Luciferase. The cell-free culture medium was used as a blank control.

### Determination of the Transgene Copy Number by Droplet Digital PCR

Droplet digital PCR (ddPCR) was used to select PGC clones with a single genomic transgene integration and was performed as described previously with slight modifications (46). Briefly, 500ng of gDNA was digested with 20 units XbaI (New England Biolabs, Germany) for one h, followed by an inactivation step at 65°C for 20 min. The Taqman PCR reaction was set up using 10ng digested DNA, 2× ddPCR supermix for probes (no dUTP) final concentration 1× (Bio-Rad Laboratories, USA), a 20× target primer/FAM-labeled probe mix, and a 20× reference primer/HEX-labeled probe mix, which was followed by droplet generation using the QX200 Droplet Generator by mixing 20ul of the TaqMan PCR reaction with 70ul droplet generator oil in a DG8 Cartridge. The cycling conditions comprised a 95°C for 10min, followed by 40 cycles of 94°C for 30 sec, 59°C for 60 sec, and a final hold for 98°C for 10 min with a 2°C/s ramp rate at all steps. The copy number was determined by calculating the proportion of positive and negative droplets using a QX200 droplet reader, which was then analyzed using Quantasoft software (Bio-Rad Laboratories, USA) (Supplementary Figure 1). The hygromycin fluorescent labeled-probe ([5′FAM-TCGTGCACGCGGATTTCGGCTCCAA-3′] along with the primers: ddHygro_F [5′-CATATGGCGCGATTGCTGATC-3′] and ddHygro_R [5′-GTCAATGACCGCTGTTATGC-3′]). As a reference gene, we used the beta-actin probe ([5′HEX-GTGGGTGGAGGAGGCTGAGC-BHQ3′] along with the primer combination ddBeta_actin_F: [5′-CAGGATGCAGAAGGAGATCA-3′] and ddBeta_actin_R: [5′-TCC ACCACTAAGACAAAGCA-3′]). The quantification was done using the QX100 system (Bio-Rad Laboratories, USA) (Supplementary Figure 1).

### Stimulation of duck *RIG-I* expression in PGC-derived fibroblasts

The generated *RIG-I*-expressing PGCs were differentiated into fibroblasts (PGCFs) as previously described (47) and subsequently infected with a LPAIV H9N2 to examine the ability of *RIG-I* to detect its ligand. Briefly, 50.000 cells were seeded into 48 well plates and infected with an MOI of 0.1 for 18h before they were fixed and stained using immunofluorescence, as previously described (48). Cells were fixed with 4% PFA and kept on ice for 10 min. Subsequently, they were washed with PBS and permeabilized with 0.5% Triton X. The FLAG-Tag was detected by using a mouse anti-FLAG antibody (Supplementary Table 1) that was incubated for 1h, followed by staining with goat anti-mouse Alexa 568 (Supplementary Table 1). Slides were subsequently covered with a mounting medium that contains DAPI for staining the nucleus and covered with coverslips (Vector Laboratories, Inc., USA). Fluorescence microscopy was performed with a fluorescence microscope (ApoTome, Zeiss).

### Generation of *RIG-I*-and *RNF135*-expressing chickens

White Leghorn layer chickens (Lohmann selected White Leghorn (LSL), Lohmann-Tierzucht GmbH, Cuxhaven, Germany) were used to generate transgenic chicken lines. Animal experiments were approved by the government of Upper Bavaria, Germany (ROB-55.2-2532.Vet_02-18-9). Experiments were performed according to the German Welfare Act and the European Union Normative for Care and Use of Experimental Animals. All animals received a commercial standard diet and water *ad libitum*. PGCs that express either *RIG-I* or *RNF135* were generated using the DNA constructs shown in Fig. 1b, as previously described (49, 50). To ensure the stable integration of the transgene, we used the phiC31 integrase-mediated integration (51). Briefly, LSL PGCs were derived from the blood of male embryonic vasculature at stages 13-15, according to Hamburger and Hamilton (52). They were cultured at 37°C in a 5% CO2 environment using modified KO-DMEM as described previously (53). A total of 5×10^6^ cells per transgene were washed with phosphate-buffered saline (PBS) and resuspended in 100µl Nucleofector^TM^ Solution V (Lonza, Germany) containing the expression vector (Fig. 1b) and the integrase expression construct. Electroporation was performed using an ECM 830 Square Wave Electroporation System (BTX, USA), applying eight square wave pulses (350V, 100µsec).

After clonal selection using puromycin for *RIG-I* and blasticidin for *RNF135*, PGCs were genotyped and tested for clones with one single genomic integration, which were then used to generate the chimeric roosters. A total of 3000 cells with the desired genetic modification were injected into the vasculature of 65h old embryos, transferred into a surrogate eggshell, and incubated until the hatch. Upon sexual maturity, sperm was collected from chimeric roosters for genotyping. (50, 54). The germline-positive roosters were bred with wild-type hens to obtain heterozygous animals (the germline transmission rate is presented in Supplementary Table 2).

The examination of the inheritance of duck *RIG-I* and the genotyping of the offspring was done via PCR using FIREPol DNA Polymerase (Solis Biodyne) using the primer combination 613_RIG-I-for (5’-CCTAGGAGAAGCATTCAAGGAG-3’) and 563_RIG-I_Rev (5’-CTAAAATGGTGGGTACAAGTTGGAC-3’). The following PCR conditions were used: initial denaturation at 95 °C 3 min, followed by 40 cycles of 95 °C 30 sec, 60 °C 30 sec, 72 °C 20 sec, and a final extension step of 72 °C 5 min, resulting in a fragment of 298bp. The inheritance of the duck *RNF135* was examined using the primers combination 1121_RNF_For (5’GCATGGGATCAACCGACAGCATC-3’) and 1017_RNF_rev (5’CCACACACCAACTTGACTCGGTC-3’, using the following PCR conditions 95 °C 3 min, followed by 40 cycles of 95 °C 30 sec, 60 °C 30 sec, 72 °C for 1 min and a final extension step of 72 °C 5 min, resulting in an amplicon of 931bp.

Following the generation of different transgenic lines, they were monitored till sexual maturity for possible harmful phenotypes that could been reflected in weight gain or the ability to produce sperm or eggs. In addition, the immunophenotype of the generated birds was assessed at 2, 5, 8, and 12 weeks after hatch by flow cytometry.

### Isolation, culture, and infection of chicken embryonic fibroblasts (CEFs)

CEFs were isolated from 10-day-old (ED10) embryos according to the protocol published elsewhere (55). Before the isolation of CEFs, embryos were genotyped by collecting blood at ED10 and preparing a window of 0.5 cm^2^ in the eggshell that allowed access to the embryonic vasculature. CEFs were cultured using Iscove’s liquid medium containing stable glutamine (Biochrom, Germany) that was supplemented with 5% fetal bovine serum (FBS) Superior (Biochrom, Germany), 2% chicken serum (ThermoFisher Scientific, USA) and 1% Penicillin-Streptomycin-Solution (Penicillin 10,000 U/ml and Streptomycin 10 mg/ml) (Biochrom. Germany). Subsequently, CEFs were incubated at 40°C in a 5% CO2 atmosphere until infection. The infection of CEFs with an H9N2 virus (A/chicken/Saudi Arabia/CP7/1998) was done after overnight seeding 250.000 cells/well in 6 well plates and infecting them in three independent experiments with multiplicities of infection (MOIs) of 0.1 and 1. Supernatants were collected 24h post-infection from the infected cells and titrated on MDCK (kindly provided by Prof. Silke Rautenschlein, University of Veterinary Medicine, Hannover). The virus titration was done as previously described (48). An additional *in vitro* infection experiment with CEFs was conducted using the H1N1-WSN strain (A/WSN/1933), kindly provided by Prof. Bernd Kaspers, Ludwig Maximilian University of Munich. Briefly, cells of the four genotypes were seeded overnight and infected with two different MOIs, 0.001 and 0.01. 40h later, they were fixed and stained for plaque formation following the standard titration protocol(48).

### *In ovo* infection of embryonated eggs

Fertilized eggs from a crossing of *RIG-I* (+/-) and *RNF135*(+/-) were collected and incubated till ED10 or ED14. In a blind study, eggs were infected randomly with 10^3^ FFU of the H9N2 virus, as previously described (56). 24h post-infection, the allantois fluid and muscle tissue were collected respectively for viral titration and genotyping. The virus titration was done on MDCK cells as previously described (48).

### RT-PCR for the detection of transgene expression in different tissues

RNA from chicken organs, including lung, trachea, heart, liver, duodenum, thymus, bursa, and brain, was isolated with Reliaprep RNA Tissue Miniprep System according to manufacturer instructions (Promega), followed by cDNA synthesis using GoScript Reverse transcription mix (Promega). The detection of *RIG-I*, as well as *RNF135* from various tissues, was done using the following primers. *RIG-I*: 613_RIG-I-for (5’-CCTAGGAGAAGCATTCAAGGAG-3’) and 563_RIG-I_Rev 5’-(CTAAAATGGTGGGTACAAGTTGGAC-3’), while *RNF135* was detected using the following primers: 898_duRNF_for (5’-CTTGAGAGAGGTGGAGGGAGC-3’) and 899_duRNF_rev (5’-GGGCTGGTGGGAATTGTTGAGG-3’). *RIG-I* and *RNF135* PCRs produced an amplicon of 298bp and 148bp, respectively. β-actin mRNA was detected with primers Beta_actin_F (5′-TACCACAATGTACCCTGGC-3′) and Beta_actin_R (5′-CTCGTCTTGTTTTATGCGC-3′) (57), resulting in a 300-bp amplicon. The PCR was performed using FIREPol DNA Polymerase (Solis Bio-dyne) according to the manufacturer’s instructions. The following PCR conditions were used: initial denaturation at 98 °C 30 sec, followed by 40 cycles of 98 °C 10 sec, 59 °C 30 sec, 72 °C 30 sec, and a final extension step of 72 °C 2 min.

### Metanalysis of *RIG-I* and *RNF135* expression in duck tissues

A metanalysis of publicly available RNA-seq data was performed to estimate the expression of *RIG-I* and *RNF135* in duck tissues (reads per kilobase per million mapped reads; RPKM), as previously described (58).

### Enzyme-linked immunosorbent assay (ELISA)

ELISA was done to measure the total plasma IgM and IgY concentrations. Briefly, 96 well plates were coated overnight with anti-chicken antibodies IgM and IgY at a concentration of 2μg/mL (Supplementary Table 1). The next day, plates were washed three times with washing buffer, followed by a blocking step with 4% skim milk for one hour. The prediluted plasma samples in 1:3 serial dilution were pipetted in the plates and incubated for one hour. The detection was done using secondary HRP-conjugated antibodies at the concentrations mentioned in Tabe1, which were incubated for 1h at RT. This was followed by adding 100µl/well of TMB (3,3’, 5,5;-tetramethylbenzidine) substrate solution for 10 min, which was stopped by 50µl per well of 1M sulfuric acid solution. The optical density (OD) was measured using FluoStar Omega via the measuring filter 450nm and the reference filter 620 nm (Version 5.70 R2 BMG LABTECH, Ortenberg, Germany).

### Flow Cytometry

Peripheral blood mononuclear cells (PBMCs) were isolated using Histopaque®-1077 density gradient centrifugation (Sigma, Taufkirchen, Germany) and analyzed using flow cytometry. Extracellular staining was carried out to detect various chicken immune cell markers including T cell subpopulations, B cells and Monocytes (list of antibodies is mentioned in Supplementary Table 1). Briefly, 5×10^6^ cells were washed with 2% BSA diluted in PBS (FLUO-Buffer). To determine the living cell population, cells were incubated with Fixable Viability Dye eFluor 780 (eBioscience, Thermo Fisher Scientific, USA). After washing with FLUO-Buffer, primary antibodies (concentration shown in Supplementary Table 1) were applied for 20 min. Cells were washed in FLUO-Buffer to remove unbound antibodies and incubated with conjugated secondary antibodies for 20 min. Subsequently, cells were again washed and analyzed using an Attune flow cytometer (Thermo Fisher Scientific, USA). The obtained data were then analyzed with FlowJo 10.8.1 software (FlowJo, Ashland, USA). An example of the gating strategy used in this study is presented in Supplementary Figure 12.

### Assessment of *in vitro* T cell activation

PBMCs were isolated from 12-weeks-old *RIG-I* or WT birds and resuspended in RPMI medium supplemented with 10%FBS and 1% Penicillin/Streptomycin and distributed on a 48 well plate with a total of 5×10^6^ cells/well. Cells were subsequently stimulated with Concanavalin A (Con A) (eBioscience™) at a concentration of 25 μg per well. They were monitored every 24h for T cell activation via flow cytometry. Activated T cells were stained for surface expression of γδTCR1, CD25, or αβTCR2&3 using antibodies listed in Supplementary Table 1.

### Challenge infection experiment with H7N1

The challenge infection experiment was performed at the INRAE-PFIE platform (INRAE Centre Val de Loire, Nouzilly, France) and approved by the local Ethics Committee Val de Loire and the Ministère de l’ Enseignement Supérieur et de la Recherche under the number 2021120115599580. Transgenic birds required for the experiment were hatched in dedicated hatching isolators and genotyped by PCR using DNA extracted from EDTA blood samples collected at day 7 post hatch. At 3 weeks of age, transgenic and WT-chickens were distributed into four different groups (Supplementary Table 3) corresponding to four BSL-3 isolator units. Two other groups were kept together as MOCK controls in a single isolator (Supplementary Table 3). The birds were fed a commercial standard diet and provided water ad libitum throughout the experiment. Prior to infection (0 dpi), body weights were recorded, and blood samples were taken by occipital sinus puncture. The virus used for challenge infection was A/ Turkey/Italy/977/1999 (kindly provided by Dr. Ilaria Capua, Istituto Zooprofilattico Sperimentale Delle Venezie, Legnaro, Italy), an H7N1 subtype virus that, despite its classification as an LPAIV, causes up to 50% mortality in experimentally infected White Leghorn chickens (24, 59, 60).

Chickens were PBS/mock-treated (groups 5 and 6) or virus-infected (groups 1-4) by intra-tracheal and intra-choanal cleft inoculation of 0.1mL PBS or 5×10^5^ EID50/1×10^6^ EID50 H7N1. Animal behavior and clinical disease signs were monitored twice daily during the trial. Clinical signs were evaluated according to the following score: 0 (no clinical signs), 1 (mild clinical signs), 2 (severe clinical signs), or 3 (dead/euthanized). Birds were euthanized by pentobarbital injection into the occipital sinus at the end of the experiment or once humane endpoints were reached. Samples collected from euthanized birds included lung, caeca, and spleen. In addition, a scoring system was used to evaluate macroscopic lung lesions as follows: 1 (mild, localized edema and fibrinous exudate), 2 (moderate edema and with hemorrhage and fibrinous exudate over ∼1/4 of the lung), or 3 (severe hemorrhage and extensive edema over ∼1/2 of the lung) (Supplementary Figure 13). Data collected from the animal experiment were assessed at 2 and 6 dpi, representing the time points when tissue samples for the Fluidigm assay were collected. Details regarding the number of birds, group distribution and number of analyzed samples are given in Supplementary Tables 3 and 4.

### Quantification of the viral genome in H7N1-challenged chickens

Total RNA was isolated from the isolated organs and conserved in 1ml NucleoProtect RNA (Macherey-Nagel, Germany). Samples were later processed for total RNA isolation with NucleoSpin RNA (Macherey-Nagel, Germany) according to the manufacturer’s instructions. The viral genome of the H7N1 virus was quantified using qRT-PCR that was conducted with the Bio-Rad iTaq™ Universal SYBR ® Green One-Step Kit (BioRad, California, USA) using primers designed for the detection of the M gene as published elsewhere (61).

### Analysis of gene expression via Fluidigm Dynamic Array

Gene expression was analyzed as previously described (25). Briefly, total RNA was extracted from infected animals’ lungs, caeca, and spleen and processed for quality control via nanodrop. Reverse transcription was performed using the High Capacity Reverse Transcription Kit (Applied Biosystems) according to manufacturer’s instructions with random hexamers and oligo (dT)18 in a final volume of 10μl, containing 250 ng total RNA. Subsequently, the cDNA was pre-amplified using TaqMan PreAmp Master Mix (Applied Biosystems). Quantitative PCR was performed in the BioMark HD instrument with the 96.96 IFC Dynamic Array (Fluidigm). The reaction was prepared by mixing 2.5 μl TaqMan Gene Expression Master Mix (Applied Biosystems), 0.25 μl 20X DNA Binding Dye Sample Loading Reagent (Fluidigm), 0.25 μl 20X EvaGreen DNA binding dye (Biotum) and 2 μl of preamplified cDNA. The qPCR was run under the following thermal conditions: 50°C for 2 min, 70°C for 30 min, 25°C for 10 min, followed by hot start 50°C for 2 min, 95°C for 10 min, PCR (x30 cycles) 95°C for 15 sec, 60°C for 60 sec and melting curve analysis 60°C for 3 sec to 95°C.

Raw quantitation cycle (Cq) data were collated with the Real-time PCR Analysis software v3.1.3 (Fluidigm), setting parameters of quality Cq threshold to auto (global) and the baseline correction to derivative. Raw Cq values were processed with GenEx.v6 MultiD, with correction for primer efficiency and reference gene normalization. Stability of the expression of reference genes: *TATA box binding protein (TBP), Tubulin alpha chain (TUBA8B), beta-actin (ACTB), beta-glucuronidase (GUSB), glyceraldehyde-3-phosphate dehydrogenase (GAPDH) and ribosomal 28S (r28S)* were evaluated via NormFinder (GenEx). The geometric mean of the most stable (*GAPDH*, *GUSB* and *TBP*) were used for normalization. Technical replicates were averaged, and relative quantification was to the maximum Cq value obtained per gene, transformed to logarithmic scale, which was then statistically analyzed using T-test.

### Challenge infection experiments with H3N1 and H9N2

Challenge experiments were performed at the Animal Research Center (ARC) of the Technical University of Munich and approved by the government of Upper Bavaria under the number ROB-55.2-2532.Vet_02-21-11. The H3N1 virus (A/Chicken/Belgium/460/2019) was kindly provided by Dr. Joris Pieter De Gussem (Poulpharm BV). Infection with H3N1 was done with 28-week-old chickens. Infection with H9N2 (A/chicken/Saudi Arabia/CP7/1998) was done with four weeks old chicks. According to the obtained genotype at hatch (WT, *RIG-I*-and *RIG-I-RNF135*-expressing chickens), birds were distributed to groups of four or six birds per group. An infectious dose of 10^6^ FFU in 0.2mL PBS per bird was distributed via nasal and tracheal routes for both viruses. In both experiments, birds were monitored daily for clinical symptoms, and tracheal swabs were collected on days 0, 3, 7, 11 and 17 to analyze the viral RNA loads by RT-qPCR(48). Duodenum, lung, and spleen samples were collected when mortality occurred or on the final day of the experiment (17 dpi). The expression of *RIG-I*, *RNF135*, *IFN-γ, IFN-α, IL1β, and IL6* was assessed by RT-qPCR (62).

### Histology

Caeca and lungs (H7N1 infection experiment) or oviduct samples (infundibulum, magnum − H3N1 infection experiment) were fixed in 10% neutral buffered formalin and processed routinely. Four-micrometer tissue sections were stained with hematoxylin and eosin for light microscopy. The sections’ analysis addressed histopathological changes and signs of inflammatory reactions, including degeneration, necrosis, infiltration of inflammatory cells, fibrin exudation, and epithelial hyperplasia.

### Statistical analysis

Statistical analysis was done using IBM SPSS Statistics (Version 28.0.1.1. IBM, Armonk, USA). The normality of the data was examined via Kolmogorov-Smirnov and Shapiro-Wilk tests. The comparison between two groups was made either with Two Samples T-test or Wilcoxon Signed Rank test. Multiple group comparison was made using Kruskal-Wallis Test or with One-Way ANOVA. The statistical test result was considered significant when the *P* value was less than 0.05.

## Acknowledgments

Animal husbandry and experimental infrastructure was provided by the TUM Animal Research Center (ARC). The authors thank the animal keepers of the ARC as well as the team at the INRAE Plateforme d’Infectiologie Expérimentale (INRAE-PFIE, Nouzilly, France) for conducting the H7N1 challenge infection experiment. Additionally, the authors thank the technical staff of the involved laboratories for excellent assistance.

## Author Contributions

H.S., T.v.H., S.Schleibinger, R.K., L.H.N., H.V., R.G., V.G., R.S., M.N.A., B.B., B. Schade, D.E., S. Sives, L.V., S.T., and B. Schusser contributed to the production of experimental data. H.S., T.v.H., S. Schleibinger, L.H.N., R.G., B.B., B. Schade, D.E., S. Sives, L.V., S.T., and B. Schusser contributed to experimental design and interpretation of the data. H.S., S.T., and B. Schusser wrote the manuscript.

