## Supporting information for "Genetic Reinstatement of RIG-I in Chickens Reveals Insights into Avian Immune Evolution and Influenza Interaction"

**This file includes:**

Figures S1 to S13
Tables S1 to S4

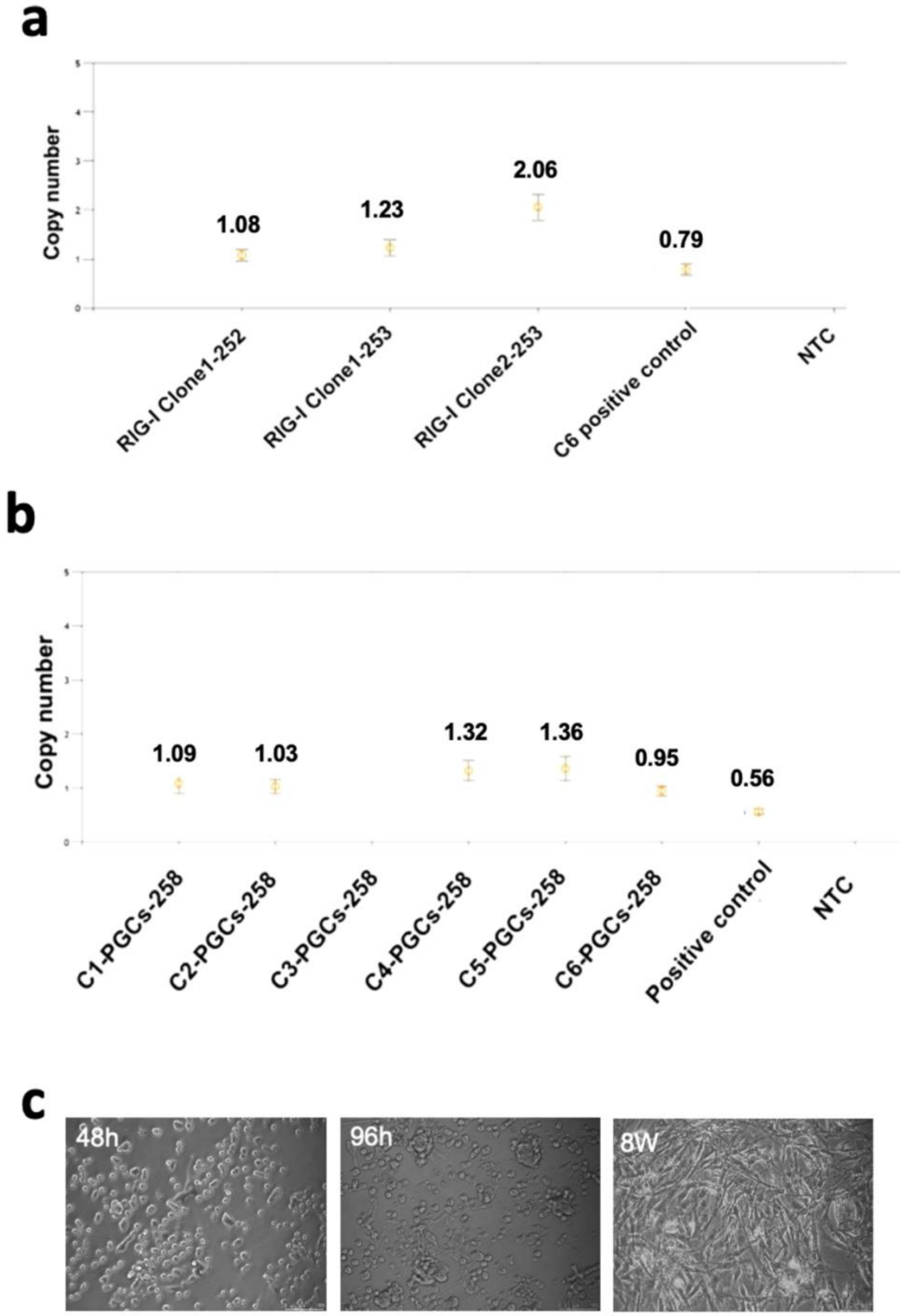


**Fig. S1. Results of Droplet Digital PCR (ddPCR) of PGCs that express the duck *RIG-I* (a) and those that express the duck *RNF135* (b)**

**a.** #253 indicates the construct with the full *RIG-I* length promoter used to express the duck *RNF135* in PGCs. Clone #1 was used for the generation of *RIG-I* chimeric roosters.

**b.** #258 indicates the construct number used to express the duck *RNF135* in PGCs. Clone #6 was used for the generation of *RNF135* chimeric roosters.

**c.** A representative figure of selected clones that were derived into fibroblasts (PGC-derived fibroblasts) and tested for the *RIG-I* activity by the detection of FLAG-Tag

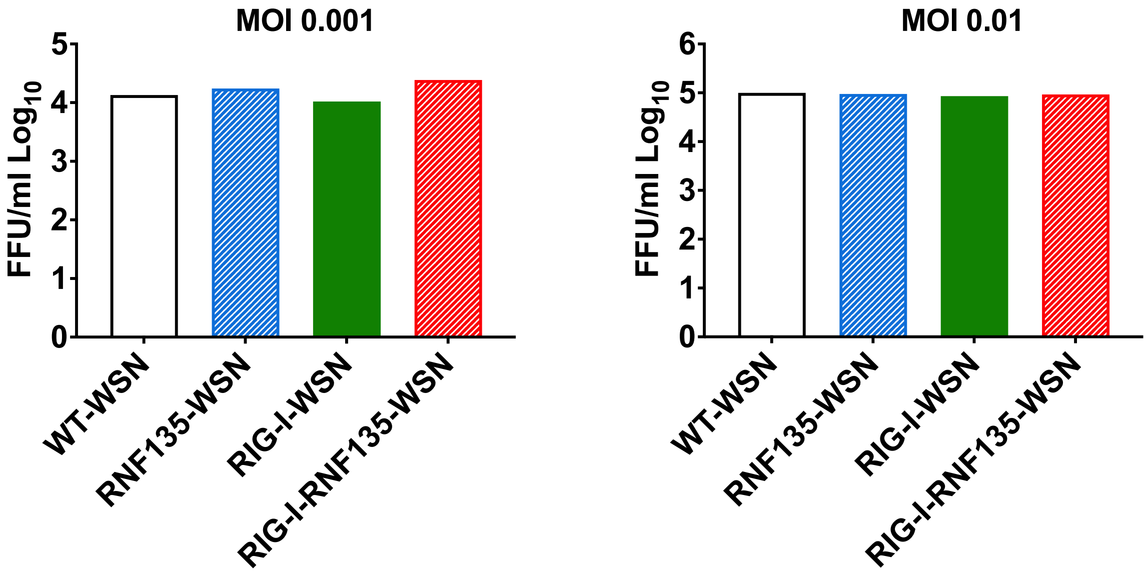


**Fig. S2.** No differences in viral replication upon infection of CEFs infected with H1N1-WSN (WSN) at two different MOIs 0.001 and 0.01. Cells were infected with WSN for 40 hours and then processed for staining of viral plaques.

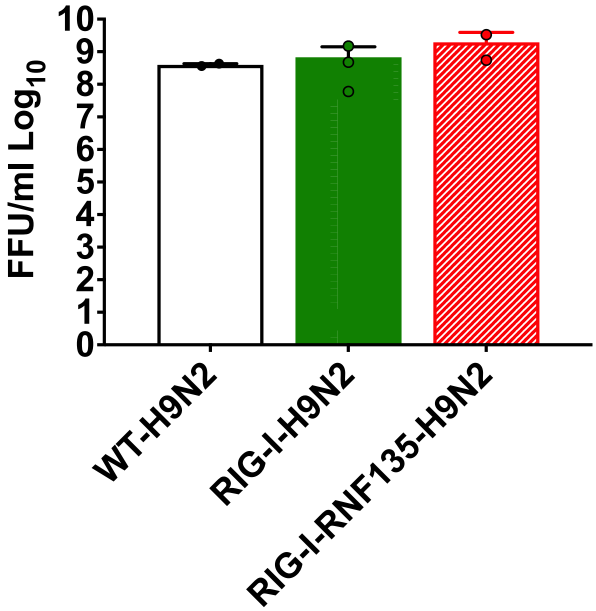


**Fig. S3.** Quantification of newly produced viral particles after infection of embryonated eggs. 14-day-old embryonated eggs were infected with LPAIV H9N2 at $10^3$ FFU/egg; Allantois fluid was collected 24hpi and titrated on MDCK cells.

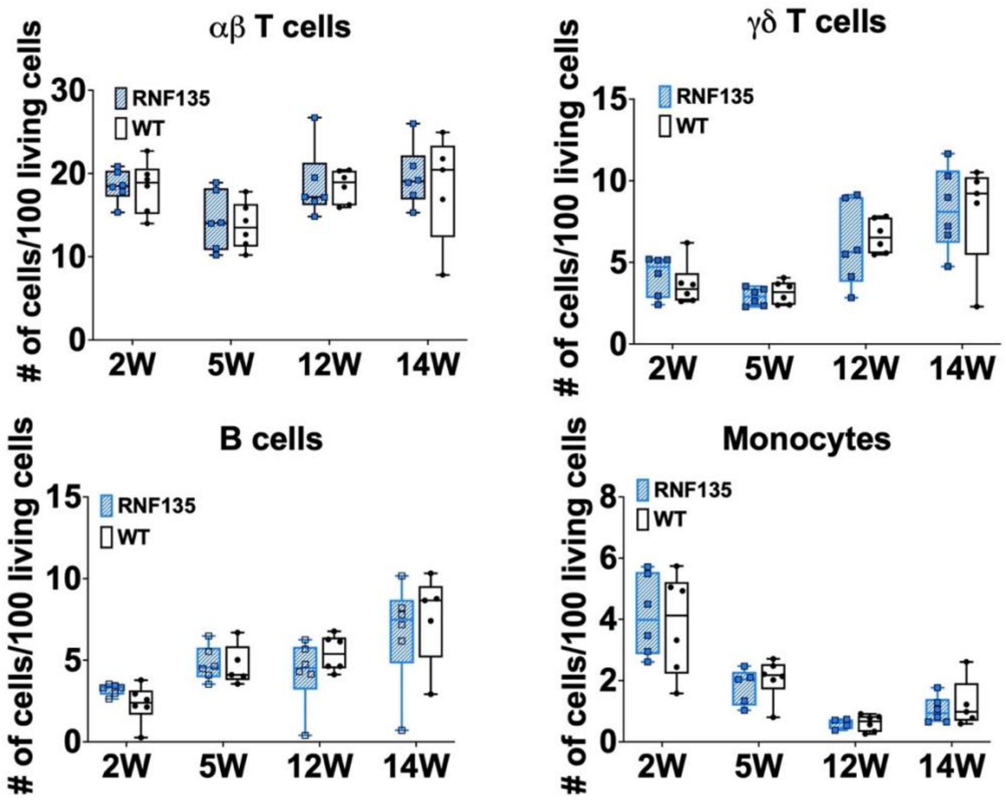


**Fig. S4.** Immunophenotype of *RNF135*-expressing chickens

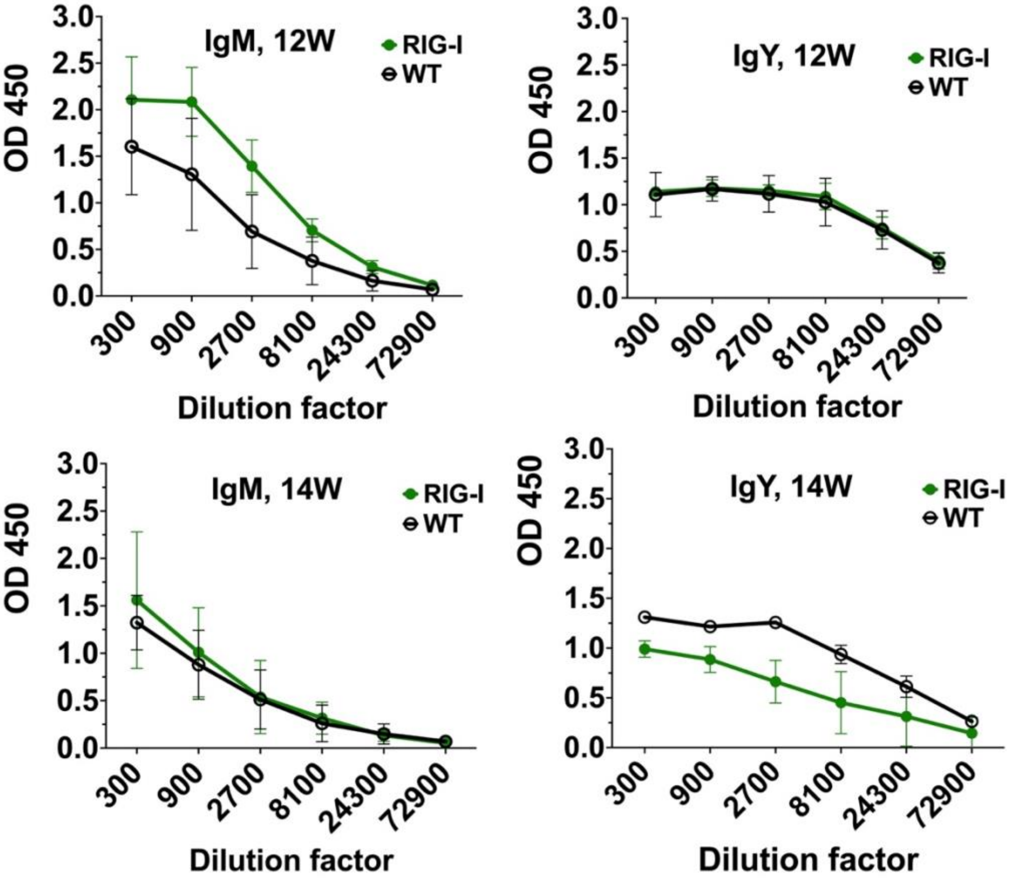


**Fig. S5. Immunoglobulin levels in *RIG-I*-expressing birds compared to WT** Relative amounts of total plasma IgM and IgY in *RIG-I*-expressing birds and their WT siblings compared by ELISA at 12 weeks of age (n≥3)

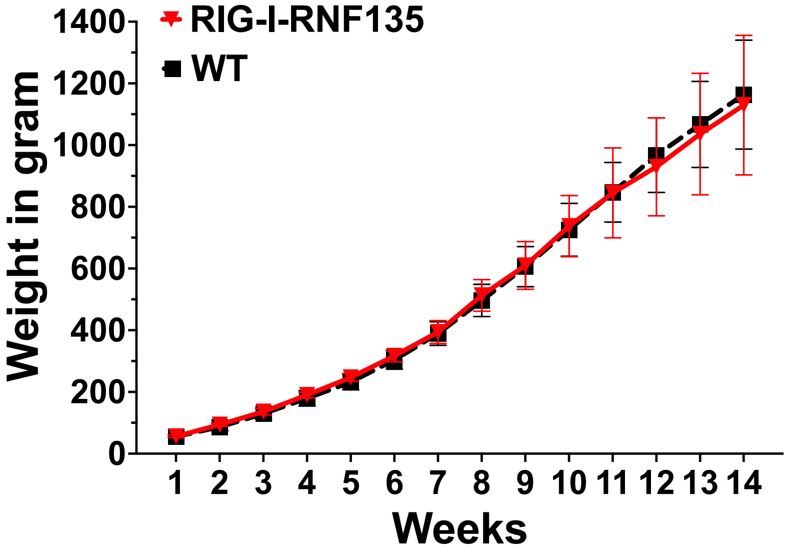


**Fig. S6.** Body weight development over 14 weeks of *RIG-I-RNF135*-expressing chickens in comparison to WT siblings (n ≥4)

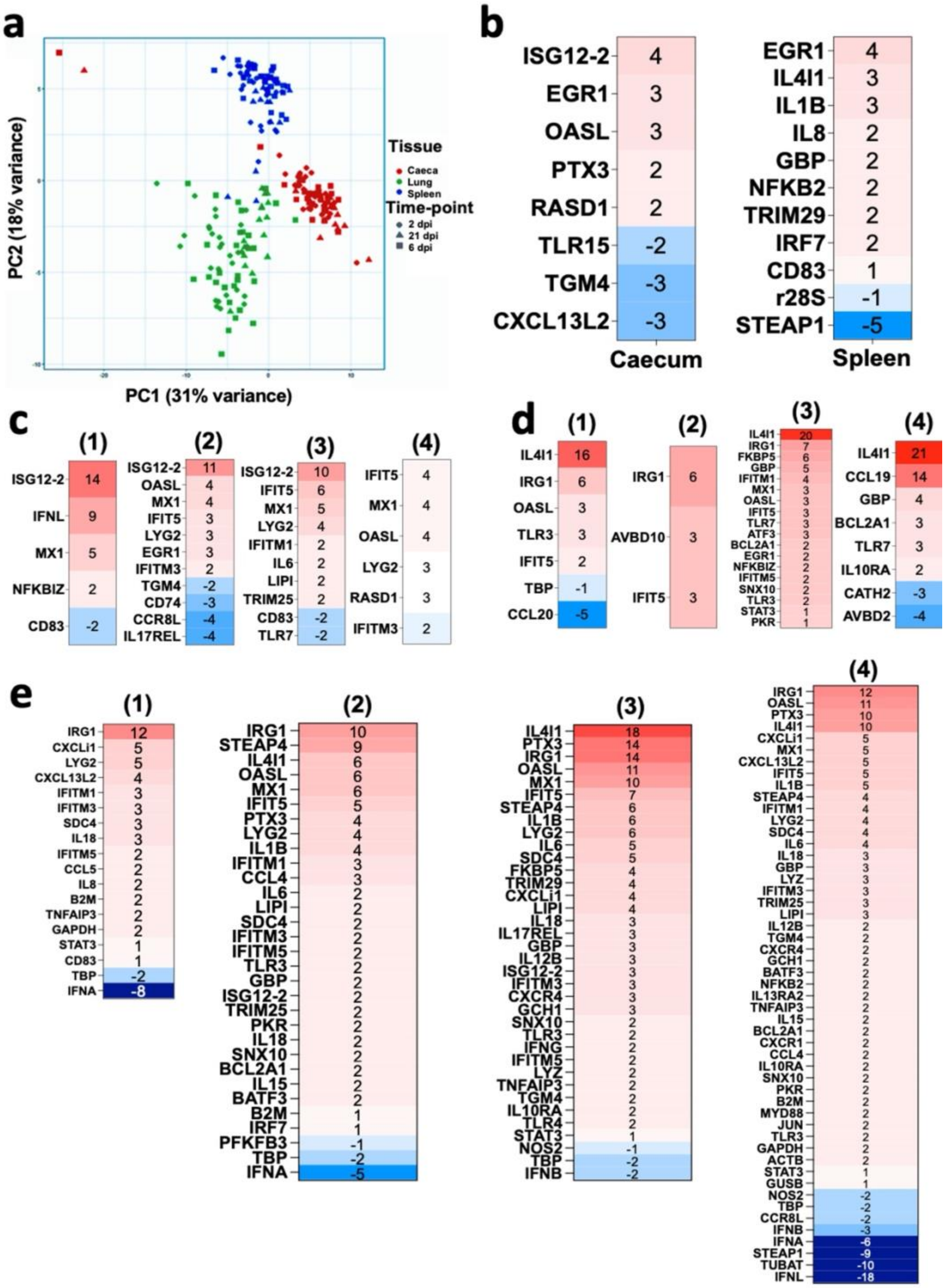


**Fig. S7. Fluidigm qRT-PCR array of naive and challenged birds with H7N1**

**a.** Principal component analysis (PCA) of global gene expression of viral innate immune-related genes. **b.** *RIG-I-RNF135* MOCK compared to WT-MOCK birds; positive values indicate high expression in the *RIG-I-RNF135* expressing chickens, while negative values indicate high expression in WT-MOCK. **c.** Expressed genes in the caecum of infected groups compared to WT-MOCK at 2dpi; positive values indicate high expression in the infected groups (1=WT-H7N1, 2=*RIG-I-RNF135*-H7N1, 3=*RIG-I*-H7N1, 4=*RNF135*-H7N1), while negative values indicate high expression in WT-MOCK. **d.** Expressed genes in the lung at 2dpi; **e.** expressed genes in the spleen at 2dpi. Significant DEGs were identified by comparing the relative expression values for every chicken line to the WT-MOCK individually per timepoint, with a significance level set at $p < 0.05$; fold change >1 (n ≥5/timepoint).

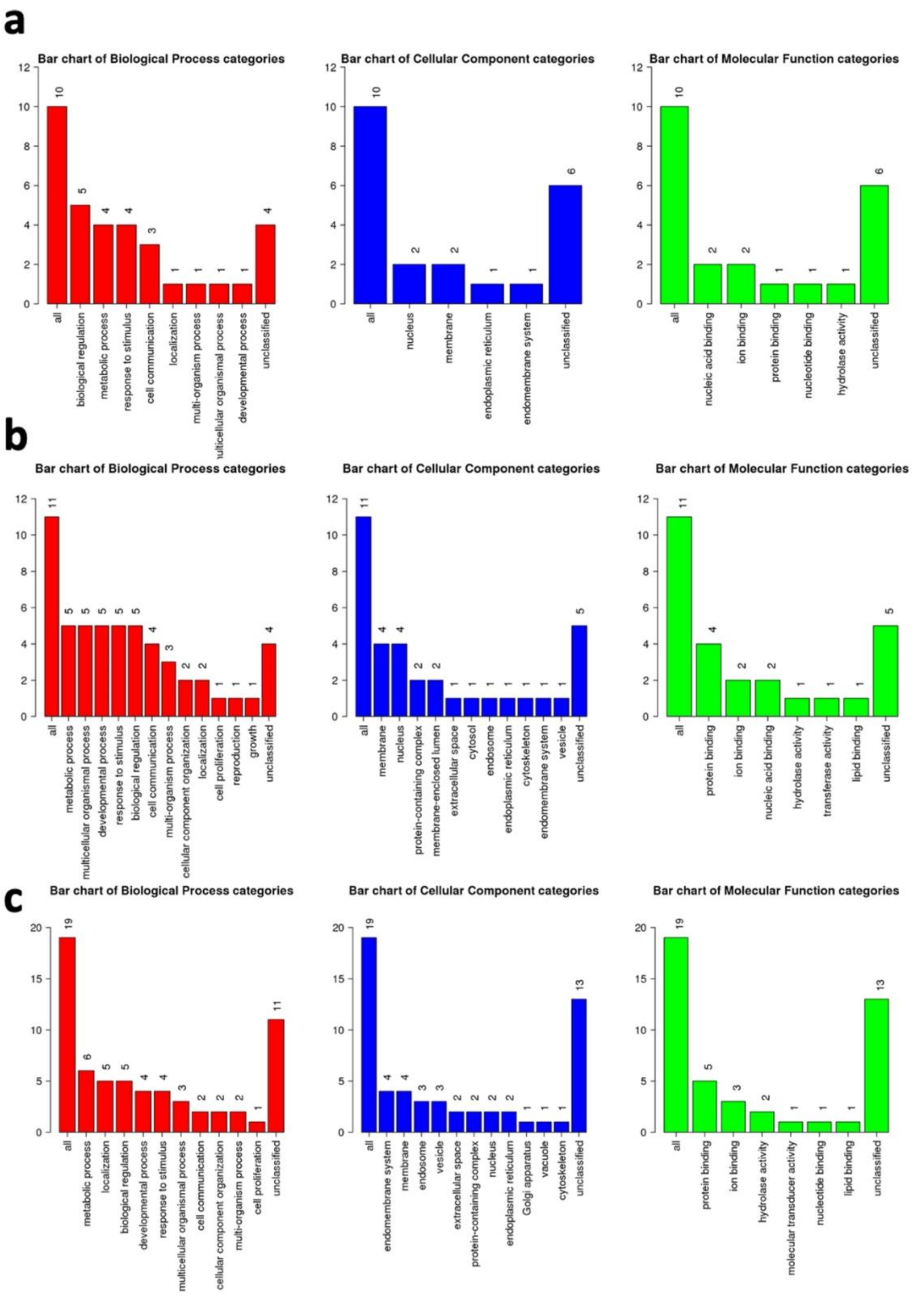


**Fig. S8. Involvement of the regulated genes in the biological processes.** Each category is represented by a different color (Biological Process = red, Cellular Component

= blue and Molecular Function = green). **a.** regulated genes in the caecum and spleen of *RIG-I-RNF135*-expresing birds. The height of the bar represents the number of IDs in the uploaded list of genes and the category. **b.** Function of genes expressed in the lung and caecum of *RIG-I*-expressing chickens **c.** Function of genes expressed in the spleen of *RIG-I*-expressing chickens at 6dpi. Figures were generated using WebGestalt (WEB-based GEne SeT AnaLysis Toolkit).

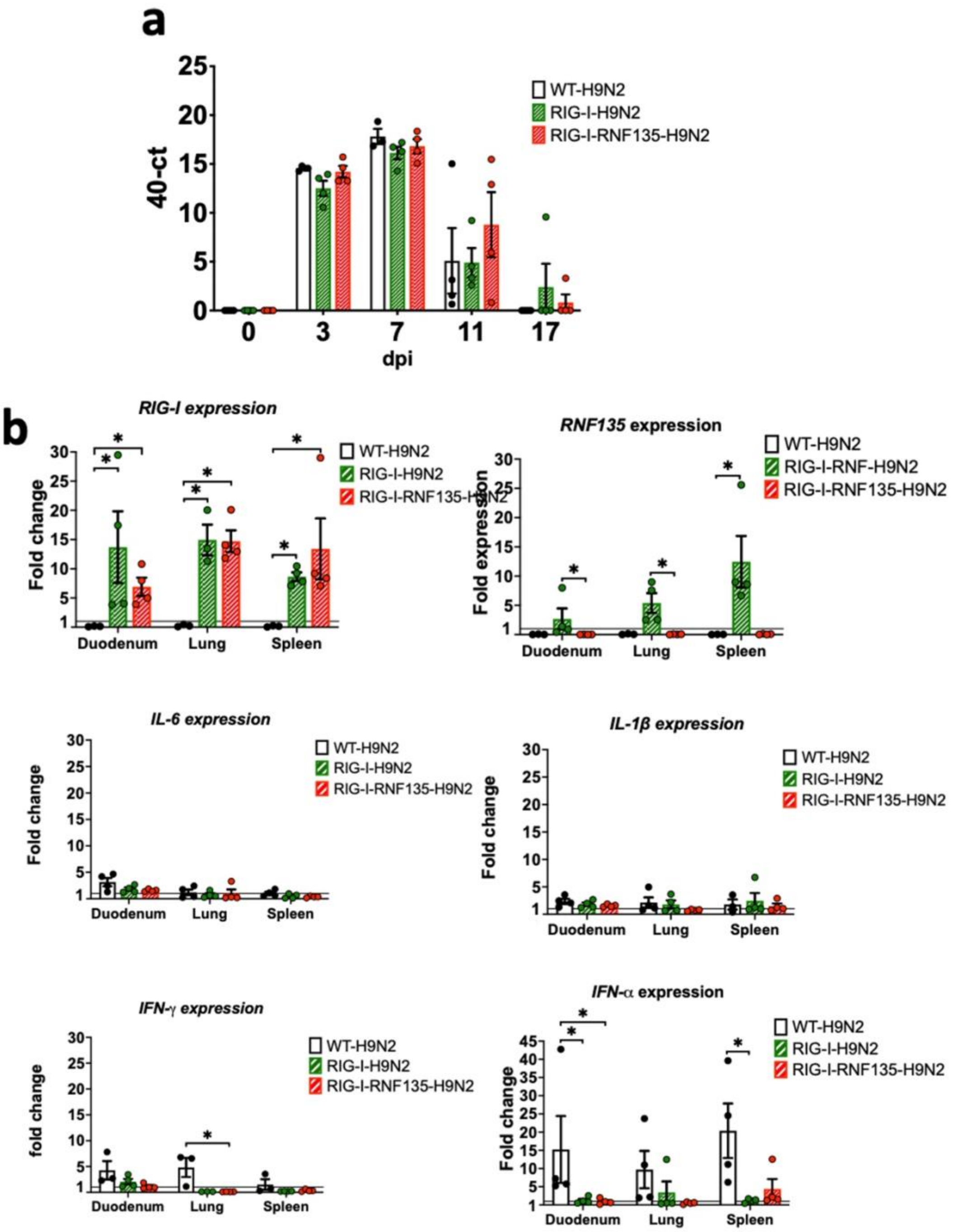


**Fig. S9. H9N2 infection does not lead to an exacerbated pro-inflammatory response in *RIG-I* and *RIG-I-RNF135*-expressing chickens:** The generated transgenic chickens were challenged at four weeks of age with H9N2 and assessed for different parameters. a. Viral shedding based on tracheal swabbing and viral RNA load analysis. b. Expression of *RIG-I*, *RNF135*, and influenza-regulated genes in the duodenum, lung and spleen. Error bars indicate standard error of mean (SEM); (*) indicate statistical differences between groups tested simultaneously ($p<0.05$). Depending on the normal distribution of the data, multiple group comparison was done either with one-way ANOVA or Independent-Samples Kruskal-Wallis Test

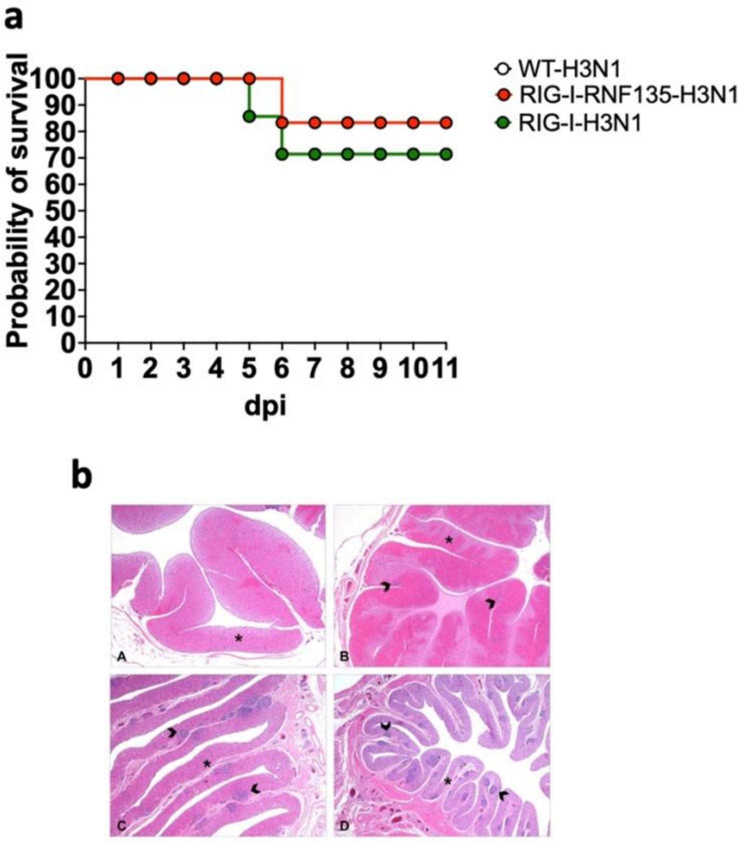


**Fig. S10. Challenge infection experiment with H3N1 a**. Probability of survival in the H3N1 challenged groups; **b.** A representative figure of histopathological scoring; A: Magnum, normal with physiological height of mucosa, B: Magnum, grade 1, mild lymphoplasmacellular salpingitis with physiological height of mucosa, C: Magnum, grade 2, moderate lymphoplasmacellular salpingitis with prominent atrophy of the mucosa, D: Magnum, grade 3, severe lymphoplasmacellular salpingitis with severe atrophy of the mucosa

Asterix: height of mucosa
Arrowhead: lymphoplasmacellular salpingitis
Hematoxylin and eosin (HE); 20x, respectively

**Duck RNF135 promoter (*Anas platyrhynchos*)**
GGAGCAGAGCCAGGCAGCTATAGTCTACCTGACTGCAGCTCCTATCCAGACAGGGACATGTTTCTTTTCCATGGAGTGGTTTTAATGCCTTTTACAGTGTTAGATTTCTG
TCACAGAAATTGTAAAAAAAATAAAAGTCAAGTTTGAATTGTGGAGAATACTGTAATGGAATCACTCTAATGACAGATGTCCCTACCTCCTTGCTTTCAACAGTCTCTGCTA
TTTGATTCTTTTATGAGTGGCAAATGAGACATGTAAAGACTTTTTTTTTCTTTTCCAAAGCAACCCCCACATACTGCCTCAGTCAAGACGAGTTCATGATGAAGACACACTG
ATTGGTATTACCTCCTAAAAACAGCTAAAGCTTTGGGTCATGAGTTCACTTACTGAATTTGAGTAGTCTGAGCAGTGAAAGGCAGAAGTCAGTTTTAAAAGTGCTGTCTTC
CTTGGCTTGTTTGAAGACACTCAATGTATGAGGACCAGCACGGAAAGCACAGAGTGAGTGCCTCCTGATCAGGTGCTACGTACATCCATTCTGCGCAGGAATGGGTGA
GGATTTGCACTGGCTCACAAAGGGCACTGAGTGTTTTCTTGCCTCACAGCCAAGCCCAAGATTTGGTTCTACCAAGAAGTCAGTCATTCAGATATTTAATTCTTCTTTTCA
GATTCCCTTAGTTTGCAAATCAAGCGGAAGTCTTGCAAACTAAGCTTCTGGTAAAGCTTCCACCTTTTGCTGTTGCTTGCCAGCCAGTCTAAGGCACCCCAAATAGTTCTA
CAGAAAGAAGGAAGTACCAATATATGCAGGGATTAGTGTGATTAACTCTTTTTTTCCCACAACAGTACAGAAAATTTGACTTGGTTCATCATTCTGTCTGTCTTAGTCAACT
CATCATCTTCCAGCCTTGAACAAACTGCAGGAGTTTTGCTTCCTGACATTCTCCCATAAACTCCCTGATGTTTCTGTTCTTTCTTTCCTCTCCCCATACGATCTGGGGAATT
ACAGTTCACGTGCTGGTTCAGCAAATCTTGTAACAGCCTCATCCTGCAGCTCCTGAAATCAACAGCGAGGCACTTGTTGAAGATCATAGTCTGGTTTGAGATGCTTTTTT
TTATTATTTATTTTGAGGTGAGCTGAGCAATTCTGGATGATCTCAAAAAGTTTAGTCTACTCAGAAGCCTCAGCTTCACGGATGGGGTTCCCTGTGTGATGTCCGGGAAA
GGTCAGCAGACCAACCGTGTCAGCAACCCTGTTACAAAACCAGGAAATTCCCAGAAGAAAAAAAAAAATGCTACTGTGCGAAGACGACAACTTTTTGCAGACTGGAGGC
GAGGTTACGGAGAGCTCCGTGTGACGGTACCTTCCTACAGCCAACCCAAGTCCTGGCGCCGAGCTGAAGGCAGCAGCAGCACCGGAGCCCGGCTTCGCGGGTGCCC
CGGGCCTTCAGCTGCCGGTAGCCCCGCACCGGGCCCGGTCCAGCACCGAGCCCAGCCCCACACCGGGCCCGGCCCCCCCGGCGGGAGGCGGCGCCCCGAGGAAA
CGAAACCGCGCCGCGCTCCGCTCCGCCCCGAGCAGGACCT

**GC-optimised duck RNF135 sequence (*Anas platyrhynchos*)**
ATGTATCCTTATGATGTTCCCGATTATGCTGCGGTTCCCGCTGAGCTTGGCCGGCTCCTCGCTGACGTTGAACTCAGCTGTTCATGCTGTTTGCAGTACTTTACCGAGCC
CGTACGGTTGGCGTCTTGCTCTCACAGCTTCTGTCGGAGCTGTATCGACACGTACTGTCGGGGTCGCCGGCGGGCACCCTGCCCTCTCTGCAGAGAGGATTTTGAGC
CGAAAGATTTGCGCCCGAATAGAGAACTTGCAGCCCTTGTCTCACTCATGTTGGGCGGAGGCAGAGGTGAGGGACTGGGTGCATGGGATCAACCGACAGCATCTGGT
GACGGGGCGGGCGGAGGGTGCAGCAGTGCCTGGAGACGCCCGGGGGAAAAGGAGGAGCAAATACGGGATATTAGCAAACAGCTTGAGATAACTAAGGAAACTATTA
ATGCGCTGCGGAAAGATTTGTCTAAGACGAAAGAATATACAAGCCAAATACTCTCACAAATAACAGAGGACTTTTGCTGCATGAAAGAGTATATTGAGAGGCAAGAAGAA
AATACTTTGATGTTCATCGAACAGGAACAGAGGGCGGCCCGGCAAAAGATAGTACAAACTATCCACCAGCTTTGTGTTGAAAAATACAAATTGATAGACATAAAGGCGCA
AATGGAGAAGGGGTTGGAAAGTGACGAAATGGAATGGCAGACCTCTAACCTTCTTGAGAGAGGTGGAGGGAGCCCTAGTACCATGCATAAGTTCACAATCGATGAGAA
ATTCAACGTCGTTAGGAGCGCCGTAGGAGATCTGAAAAGAAAATTGGAGATCTTGCTGCTCGAAGAGTATCCTCAACAATTCCCACCAGCCCAAAGTCCCGACTTGCAT
CAGGAAACGAGTGTCTGTTCATTGTCATCCGAAAGCGCGGCTAAGTCACCCGAACCGTCTATCTCATCCCAATTTTCACGCTGGGCTGATAACGTCACCTTTGACTTGAC
GACCGCCTATGATAGACTTGCCATCACCGATCAGAATAGAAAAGTTATGGTGTCATCTAACCCAACCTATTACGAGCCATCCCTTAAACGGTTCTGTATAAGTCAGGTTCT
TTGCAGCCAAGGCTTCAGTACCGGGTGCCACTATTGGGAGGTAATCACCAAGGACAGCGACGGTTGGGCGGTTGGCGTTGCCAGAGGAACCATCGGGAGACGGGAT
CGCTTGGGACGGACCGAGTCAAGTTGGTGTGTGGAGTGGGTGGGACCTCAGAAGCAGCTGAGCGCCTGGCATCGCAATCAAGAGACACTGCTGCGGAACGACAAGC
CACTTAAAGTAGGCGTGTTCCTTGAACTGCAAAAGACCGTATCTTTTTATGCGATAACCGATCGCGAGATGCTCTTGCATACATTCGAAATCAATAACAGTAACCCTTTGT
ACCCGGCCTTCTGGCTCTATAGCTTGGACAAGAACGGCTCTCTTACGATAAATCATATTAACAGGAAATAAG

**Protein sequence of the duck RNF135 (*Anas platyrhynchos*)**
M**YPYDVPDYAA**VPAELGRLLADVELSCSCCLQYFTEPVRLASCSHSFCRSCIDTYCRGRRRAPCPLCREDFEPKDLRPNRELAALVSLMLGGGRGEGLGAWDQPTASGDGA
GGGCSSAWRRPGEKEEQIRDISKQLEITKETINALRKDLSKTKEYTSQILSQITEDFCCMKEYIERQEENTLMFIEQEQRAARQKIVQTIHQLCVEKYKLIDIKAQMEKGLESDEME
WQTSNLLERGGGSPSTMHKFTIDEKFNVVRSAVGDLKRKLEILLLEEYPQQFPPAQSPDLHQETSVCSLSSESAAKSPEPSISSQFSRWADNVTFDLTTAYDRLAITDQNRKVM
VSSNPTYYEPSLKRFCISQVLCSQGFSTGCHYWEVITKDSDGWAVGVARGTIGRRDRLGRTESSWCVEWVGPQKQLSAWHRNQETLLRNDKPLKVGVFLELQKTVSFYAITD
REMLLHTFEINNSNPLYPAFWLYSLDKNGSLTINHINRK*

**Blue colored letters indicate the inserted HA tag**

**Fig. S11.** Sequence of duck *RNF135* promoter and the duck *RNF135*

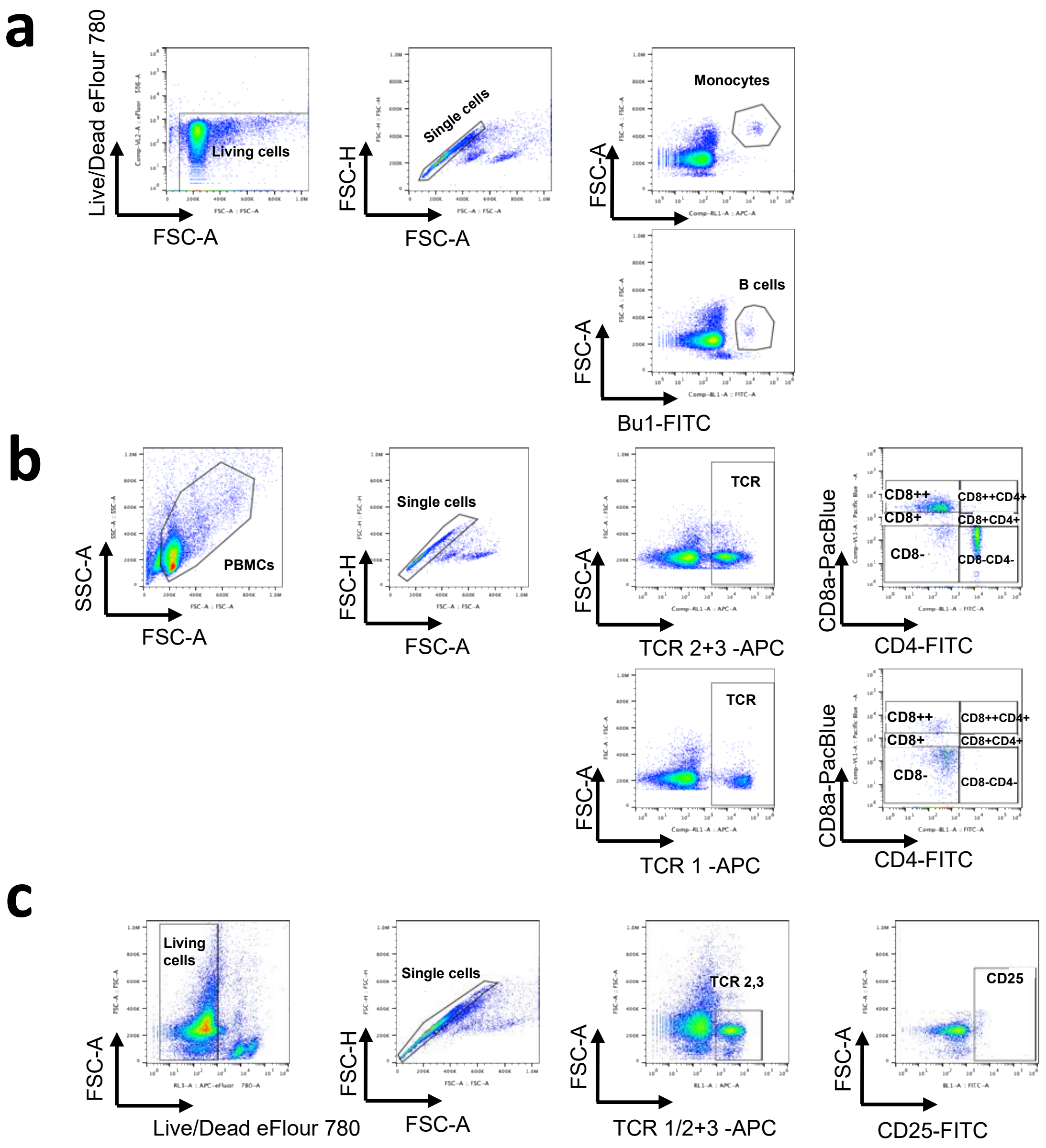


**Fig. S12. Gating strategies for the detection of different immune cells. a.** Detection of B cells and monocytes. **b.** Detection of T cell subpopulations. **c.** Live/Dead staining and detection of TCRαβ1, αβ2/CD25+ T cells and TCRγδ/CD25+ T cells

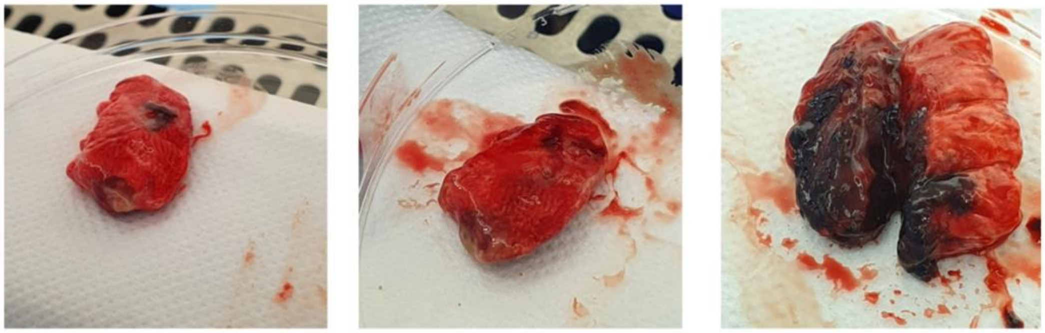

Score 1 (+)
Mild, localized edema and fibrinous exudate

Score 2 (++)
Moderate edema with hemorrhage and fibrinous exudate over about 1/4 of the lung

Score 3 (+++)
Severe hemorrhage and extensive edema over about 1/2 of the lung

**Fig. S13.** Scoring system used to evaluate H7N1-induced lung lesions.

**Table S1: Clones and concentrations of the used antibodies**

| Antibody | Company | Clone | Assay | Concentration |
|---|---|---|---|---|
| Mouse Anti-Chicken TCRγδ-UNLB | Biozol | TCR1 | Flow cytometry | 0.625 µg/mL |
| mouse anti-chicken TCRαβ/Vb1-UNLB | Biozol | TCR2 | Flow cytometry | 2.5µg/mL |
| mouse anti-chicken TCRαβ/Vb2-UNLB | Biozol | TCR3 | Flow cytometry | 2.5µg/mL |
| mouse-IgG1_anti-chicken chB6_UNLAB | Biozol | AV20 | Flow cytometry | 2.5µg/mL |
| mouse IgG2a -anti-chicken KUL01_UNLAB | Biozol | KUL01 | Flow cytometry | 0.625 µg/mL |
| goat anti-mouse IgG (H+L)-APC | Biozol | Polyclonal | Flow cytometry | 0.625 µg/mL |
| mouse-IgG1_anti-chicken CD8α_PacBlue | Biozol | CT-8 | Flow cytometry | 0.625 µg/mL |
| mouse IgG2a -anti-chicken CD8ß_UNLAB | Biozol | EP42 | Flow cytometry | 0.625 µg/mL |
| mouse IgG1-anti-chicken CD4_FITC | Biozol | CT-4 | Flow cytometry | 0.625 µg/mL |
| mouse IgG1 Anti-Chicken TCRγδ-BIOT | Biozol | TCR-1 | Flow cytometry | 0.625 µg/mL |
| mouse anti-chicken TCRαβ/Vb1-BIOT | Biozol | TCR-2 | Flow cytometry | 2.5µg/mL |
| mouse anti-chicken TCRαβ/Vb2-BIOT | Biozol | TCR-3 | Flow cytometry | 2.5µg/mL |
| mouse anti-chicken Bu1-FITC | Biozol | AV-20 | Flow cytometry | 2.5µg/mL |
| mouse anti- chicken MRC1L-B | Biozol | KUL01 | Flow cytometry | 0.625 µg/mL |
| rat-anti-mouse IgG2a_PE | Biozol | SB84a | Flow cytometry | 0.125µg/mL |
| Streptavidin_APC | VWR | - | Flow cytometry | 0.2µg/mL |
| goat anti-mouse IgG (H+L)-APC | Biozol | polyclonal | Flow cytometery | 0.625 µg/mL |
| Human anti-Chicken CD25 | Biorad | monoclonal | Flow cytometery (T cell activation) | 2.5µg/mL |
| Fixable Viability Dye eFluor 780 | eBioscience | | Flow cytometery | 1:1000 |
| Goat anti-chicken IgM | Biomol | polyclonal | ELISA | 2µg/mL |
| Rabbit anti-chicken IgY (used for coating ELISA plates) | Jackson | polyclonal | ELISA | 2µg/mL |
| Goat anti-chicken IgM-HRP | Biomol | polyclonal | ELISA | 0.05ng/mL |
| Rabbit Anti-Chicken IgY-HRP (used for detection) | Jackson | polyclonal | ELISA | 0.02µg/mL |

**Table S2: Germline transmission of the injected PGC clones**

| Injected PGC clone | Germline transmission (%) |
|---|---|
| *RIG-I* | 10.2 |
| *RNF135* | 6 |

**Table S3: Group distribution and number of birds used in the experimental challenge with H7N1**

| **Genotype** | **RIG-I-RNF135** | | **WT** | | **RIG-I** | | **RNF135** | | **WT** | | **RIG-I-RNF135** | |
|---|---|---|---|---|---|---|---|---|---|---|---|---|
| **Inoculation** | **H7N1** | | **H7N1** | | **H7N1** | | **H7N1** | | **MOCK** | | **MOCK** | |
| **Number** | **(n=16)** | | **(n=16)** | | **(n=18)** | | **(n=18)** | | **(n=9)** | | **(n=6)** | |
| animal # (sex) | 204 | male | 201 | female | 202 | male | 205 | male | 210 | male | 242 | male |
| animal # (sex) | 206 | female | 209 | male | 208 | female | 216 | female | 233 | male | 248 | female |
| animal # (sex) | 213 | male | 214 | female | 211 | male | 220 | male | 252 | female | 260 | male |
| animal # (sex) | 227 | male | 215 | female | 218 | male | 235 | female | 255 | male | 268 | female |
| animal # (sex) | 222 | female | 226 | male | 225 | male | 239 | female | 264 | female | 276 | female |
| animal # (sex) | 238 | male | 228 | male | 230 | female | 257 | male | 270 | female | 311 | male |
| animal # (sex) | 246 | female | 234 | female | 231 | female | 273 | male | 288 | female | | |
| animal # (sex) | 253 | female | 237 | female | 232 | female | 274 | female | 298 | male | | |
| animal # (sex) | 258 | female | 243 | male | 240 | male | 277 | male | 290 | female | | |
| animal # (sex) | 262 | male | 267 | female | 244 | female | 279 | female | | | | |
| animal # (sex) | 271 | male | 284 | male | 247 | female | 285 | female | | | | |
| animal # (sex) | 278 | female | 301 | female | 249 | male | 286 | female | | | | |
| animal # (sex) | 280 | female | 302 | female | 259 | female | 287 | male | | | | |
| animal # (sex) | 282 | male | 303 | male | 265 | male | 292 | female | | | | |
| animal # (sex) | 291 | female | 306 | male | 275 | female | 295 | male | | | | |
| animal # (sex) | 294 | female | 317 | male | 281 | female | 307 | male | | | | |
| animal # (sex) | | | | | 283 | male | 310 | female | | | | |
| animal # (sex) | | | | | 305 | male | 322 | male | | | | |

**Table S4: Number of sampled birds for organs at different days post-infection (dpi)**

| Group | Number of sampled birds | | |
|---|---|---|---|
| | **2dpi** | **6dpi** | **21dpi** |
| RIG-I-RNF135-H7N1 | 6 | 5 | 0 |
| WT-H7N1 | 7 | 6 | 3 |
| RIG-I-H7N1 | 7 | 6 | 4 |
| RNF135-H7N1 | 6 | 6 | 6 |
| WT-MOCK | 3 | 3 | 3 |
| RIG-I-RNF135-MOCK | 3 | 3 | 0 |
